# How cognitive and environmental constraints influence the reliability of simulated animats in groups

**DOI:** 10.1101/688598

**Authors:** Dominik Fischer, Sanaz Mostaghim, Larissa Albantakis

**Affiliations:** School of Management, Technical University of Munich; Department of Psychiatry, University of Wisconsin–Madison; Faculty of Computer Science, Otto von Guericke University of Magdeburg

**Author notes:** School of Management, Technical University of Munich, Germany (Dominik Fischer). Funding sources: L.A. receives funding from the Templeton World Charities Foundation (Grant #TWCF0196).

**Keywords:** Collective behavior, evolutionary algorithms, cognitive science, Markov brains.

## Abstract

Evolving in groups can either enhance or reduce an individual’s task performance. Still, we know little about the factors underlying group performance, which may be reduced to three major dimensions: (a) the individual’s ability to perform a task, (b) the dependency on environmental conditions, and (c) the perception of, and the reaction to, other group members. In our research, we investigated how these dimensions interrelate in simulated evolution experiments using adaptive agents equipped with Markov brains (“animats”). We evolved the animats to perform a spatial-navigation task under various evolutionary setups. The last generation of each evolution simulation was tested across modified conditions to evaluate and compare the animats’ reliability when faced with change. Moreover, the complexity of the evolved Markov brains was assessed based on measures of information integration. We found that, under the right conditions, specialized animats were as reliable as animats already evolved for the modified tasks, that interaction between animats was dependent on the environment and on the design of the animats, and that the task difficulty influenced the correlation between the performance of the animat and its brain complexity. Generally, our results suggest that the interrelation between the aforementioned dimensions is complex and their contribution to the group’s task performance, reliability, and brain complexity varies, which points to further dependencies. Still, our study reveals that balancing the group size and individual cognitive abilities prevents over-specialization and can help to evolve better reliability under unknown environmental situations.

**Author Summary:** The ability to adapt to environmental changes is an essential attribute of organisms which have had evolutionary success. We designed a simulated evolution experiment to better understand the relevant features of such organisms and the conditions under which they evolve: First, we created diverse groups of cognitive systems by evolving simulated organisms (“animats”) acting in groups on a spatial-navigation task. Second, we post-evolutionary tested the final evolved animats in new environments–not encountered before– in order to test their reliability when faced with change. Our results imply that the ability to generalize to environments with changing task demands can have complex dependencies on the cognitive design and sensor configuration of the organism itself, as well as its social or environmental conditions.

## Introduction

*Intelligence is the ability to adapt to changes*. According to this prevalent perspective, possessing general intelligence [1, 2] not only enables one to perform a task correctly under already known conditions, but also to perform well under unexpected conditions. Further, in natural environments intelligent behavior is not only dependent on the (maybe limited) intelligence of the individual organism, but also involves interactions with the social and physical environment [3–5]. In addition to the examples from the animal world, it is also true in *high-reliability organizations* (e.g., aircraft carrier or nuclear power plants) that individual behavior is interrelated with the behavior of the group members. This is necessary to be able to act correctly in case of an unforeseen event [6–8].

While it seems intuitive that there is a triangular relationship between the individual, the group, and the environment, we discovered a lack of research on how individual behavior and group behavior are interrelated and depend on spatial attributes of the environment [9]. This limits our understanding of how an individual actor evolves intelligent behavior and how its physiological abilities, the social setting, and the environment constrain this evolution. More generally, several studies have investigated intelligence and knowledge on the group level, and some have modelled groups of individuals as single agents (e.g., [10–14]). These studies have their origins in a variety of disciplines and have in common that they seek to elucidate the dynamics between group members.

To shed more light on the above-mentioned issue, we wanted to ask which conditions can promote the evolution of intelligent entities that act in organized groups and can additionally adapt to environmental changes under simplified conditions in a simulated evolution experiment. Inspired and motivated by Pinter-Wollman et al. [9], we wanted to investigate how the behavior and performance of evolved “animats” (simulated agents with cognitive abilities [15, 16]) varies in different task conditions, such as changes in the proportions of static objects, dynamic objects (group members), and individual cognitive abilities. This simulation enabled us to manipulate and observe three dimensions which might influence task performance and reliability: the group size, the animats’ physiology, and the environmental design. In this study, reliability describes the ability to perform well under manipulated task conditions that the animat had not been confronted with before.

We used a genetic algorithm to let the animats’ behavior evolve under various evolutionary setups. Specifically, the animats were controlled by *Markov brains (MBs)* [16], which consisted of computational units whose functions and connectivity were determined by the animats’ adaptive genome. The animats’ task was to navigate through a two-dimensional world composed of multiple rooms without colliding with other group members (see Fig 1). There was a small penalty for each collision and a large reward for crossing gates between rooms. After an evolution of 10,000 generations, we tested the final animats under modified task conditions modeled as: a variation in group size, the complexity of the static obstacles in the environment, and interaction rules between animats constraining fitness for the task. An animat was considered reliable if its task performance remained high across many of these test conditions.

**Fig 1.**
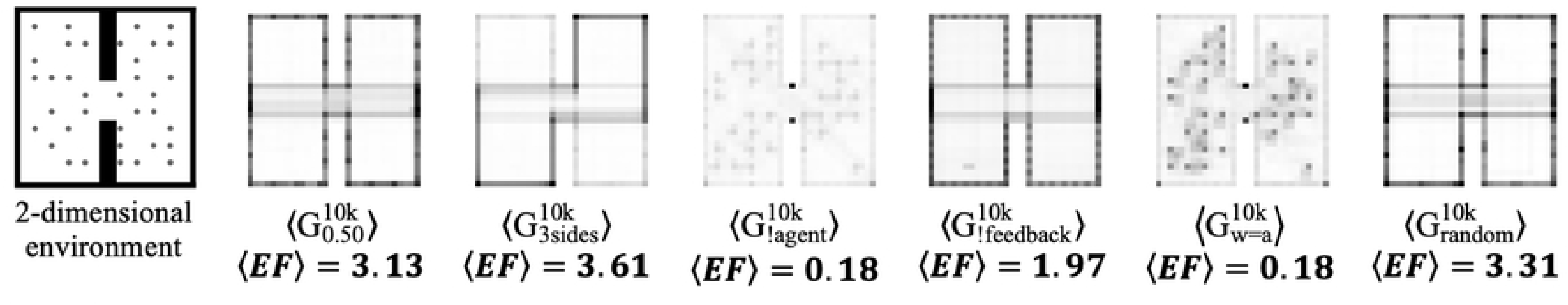
Average movement patterns of six selected conditions. The panel on the left shows the two-dimensional environment including two rooms with 36 start positions occupied (round dots). The other six panels show example movement patterns. Dark fields indicate high occupancy, and light fields indicate low occupancy in the corresponding position throughout the trial. Generally, well-performing animat groups evolve a wall following strategy. 〈EF〉 indicates the average fitness of the final generation in the specific condition.

A predecessor study focused on the influence of group size on the evolution of group fitness and reliability [17], while the present work extends the reliability experiments, includes cognitive and environmental variations in the evolutionary setup, and elaborates the measurement of brain complexity by applying measures developed within the framework of the integrated information theory (IIT) to the evolved MBs [18, 19]. There are two additional works which directly relate to our study: First, Konig et al. [20] provided the original experimental setup. They designed a two-dimensional spatial-navigation task in which a swarm of robots has to learn to travel between two rooms. Second, Albantakis et al. [19] showed how single animats evolve in a perceptual-categorization task environment with dynamic objects under various task difficulties. The primary motivation behind their work was to investigate the evolution of integrated information [18], which is an indicator for brain complexity, and its relation to task difficulty and memory capacity. In the following, we discuss how the complexity of the MBs–evolved in the various experimental setups–is related to reliability as an indicator for general intelligence.

Simulating a large set of evolutionary setups and post-evolutionary test conditions enables us to identify important cognitive and social variables and to evaluate how physical constraints influence collective movement. Specifically, the results of the simulated evolution experiments suggest the following implications: First, animats who evolve in an environment with a balanced group size evolve better reliability and can compete with specialized animats (who have already experienced changing conditions). Second, the integration of motor units into the memory network increases the performance of animats. Third, the ability to sense adjacent animats is essential for the reliability of animats to perform the task, even if it is challenging to make statements about the communication between animats in this setting. Finally, we explored how various sensor configurations influence the difficulty of dealing with the task and, therefore, the animats’ ability to cope with changes. Overall, we found that, under the right conditions, specialized animats can be reliable, that the integration of motor units has an impact on performance and reliability, that animats benefit from passive interaction, and that more sensors enable reliability with simpler and less integrated brain structures (which challenges the view that higher generalized intelligence is necessarily associated with more complex cognitive architectures). On the whole, our approach also highlights the complexity of the dependencies between the three dimensions under investigation (properties of the individual, group interaction, and environmental design), even in the simplified conditions of our simulation experiments, and thus cautions against hasty generalizations, e.g., across different species or environments.

In the following, we will first present our results on the animats’ task performance, reliability, behavior, and brain complexity across varying evolutionary setups. After that, we will discuss the findings in the broader scope of the literature and also how our work contributes to it. The last part of the work explains the methods and research design.

## Results

We simulated the evolution of artificial organisms (“animats”) with diverse cognitive architectures under various conditions for 10,000 generations (see Table 1 for an overview of all evolution simulations conducted).

**Table 1.**
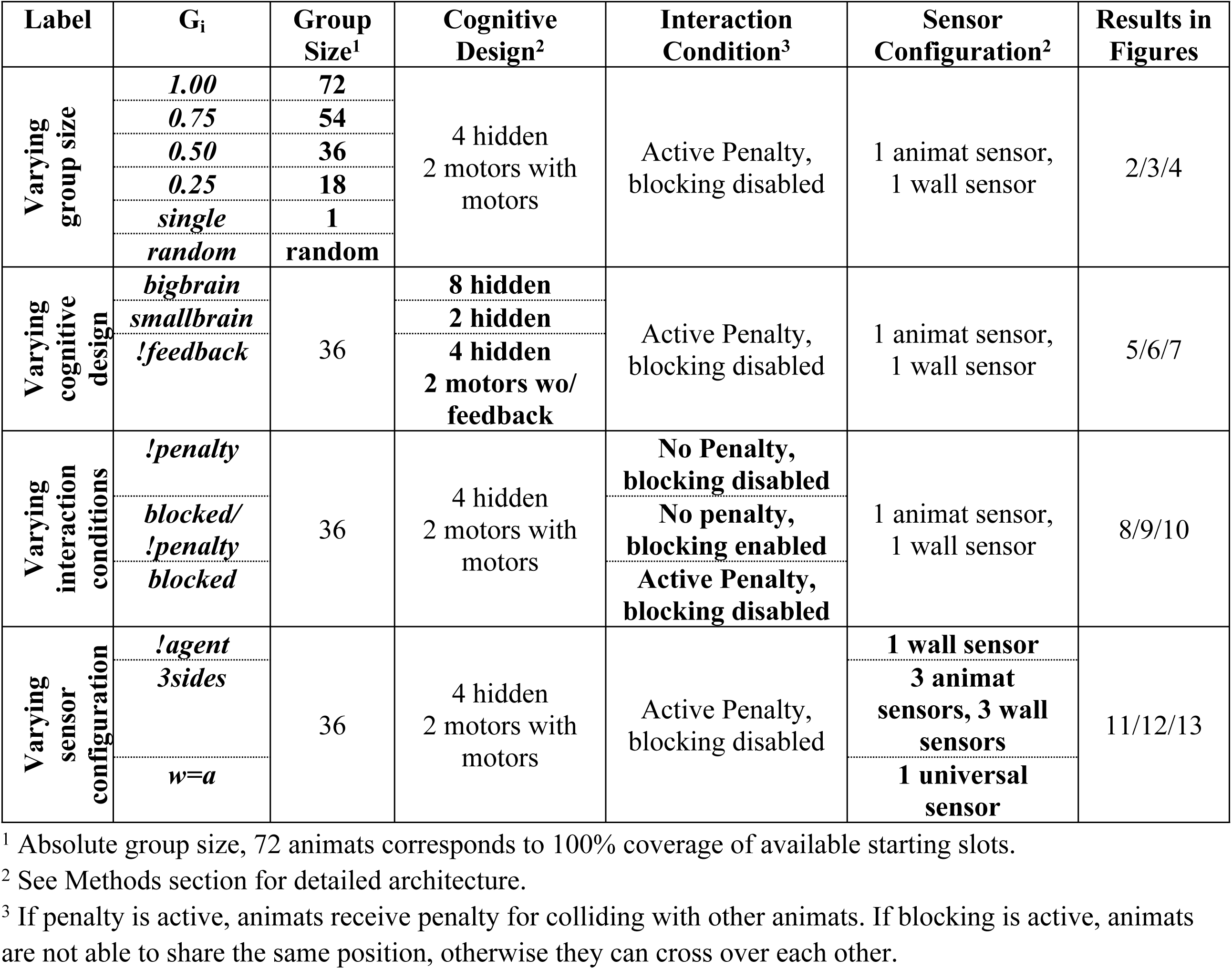
Definition of simulation conditions (“evolutionary setups”). ***G_i_*** indicates the group condition. The index ***i*** specifies the respective evolutionary setup.

All animats were evolved to travel between two rooms in a two-dimensional environment, which they shared with other animats of their same type, except in the “single” condition (see Fig 1(a) and Table 1). Fitness selection positively depended on the average number of times that the animats stepped through the gate between the two rooms. In addition, we imposed a small penalty each time they collided with other animats (if not stated otherwise). A detailed description of the task environments and the *evolutionary algorithm (EA)* is provided below in the Methods section. In many evolutionary setups, high final fitness values (***EF > 3***) was able to be achieved.

Once evolved, the final generation of animats was the basis for comparing task fitness (performance in a specific environment), behavior, and reliability (average performance across all several task environments) across conditions. In this study, we focused on assessing reliability across two dimensions: (1) the number of co-existing animats and (2) the placement of static obstacles compared to the original two-dimensional environment (see Fig 1(a), and the Methods section for details). Additionally, we varied the interaction conditions between agents as a third parameter to manipulate the agent’s reliability across group sizes.

Fig 1(b) displays six different heatmaps visualizing several evolved movement patterns. It is observable that animat groups with reasonable task fitness (***TF)*** converge towards a “swarm”-like wall-following behavior, which is driven by both interactions with fellow animats and interactions with the environment [4, 9].

We organized the presentation of our results into four sections according to the evolutionary setups thereof, as shown in Table 1 (varying “group size”, “cognitive design”, “interaction conditions”, and “sensor configuration”, respectively). Each section contains visualizations displaying the average increase in fitness across generations (“fitness evolution”), behavioral features, the reliability tests, and a complexity analysis of the evolved MBs. Since the figures are redundant in their construction, we will briefly introduce their attributes:

**Fitness:** Fig 2, Fig 5, Fig 8, and Fig 11 show (a) the fitness evolution across generations and (b) the distribution of evolved fitness values (***EF***) of the final generation. The shaded areas in (a) visualize the *standard error of the mean (SEM)* across the 30 evolution simulations that we performed per evolutionary setup. The wide bars in (b) visualize the mean evolved fitness ***〈EF〉***.

**Fig 2.**
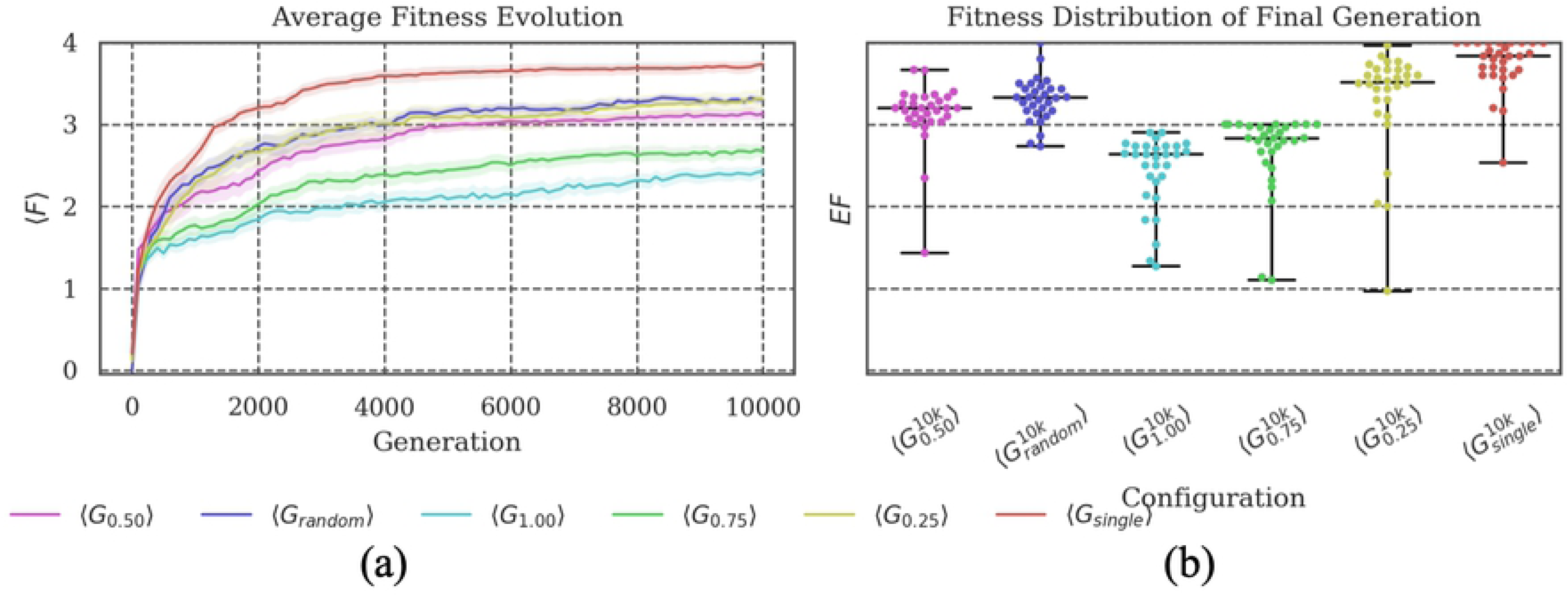
Fitness evolution and distribution of the final evolved fitness. **(a) *G_single_*** is the condition which evolves the highest fitness on average. Larger group sizes during evolution impede the animats’ fitness evolution and lead to lower final evolved fitness values. **(b)** The evolutionary setup with randomized group sizes at each generation (***G_random_***) demonstrates similar properties as those setups with fixed, intermediate group sizes (***G_0.25_*** and ***G_0.5_***).

**Reliability and behavior:** Fig 3, Fig 6, Fig 9, and Fig 12 visualize the results of testing the reliability of fitness values and behavioral features of the final generation of animats across (1) different group sizes ([***0.01389, 0.05, 0.10, …, 0.95, 1.0***]) and (2) various test conditions (changing interactions between animats and environment design). Panel (a) in Figures 3/6/9/12, shows the mean task fitness ***〈TF〉*** of testing the animats under different group sizes, respectively, in the eight different test conditions listed in Table 2. Note that the condition under which a group of animats evolved is indicated by their ***G_i_*** label (see Table 1), and ***〈TF〉*** is an average across the 30 evolution simulations per experimental setup.

**Fig 3.**
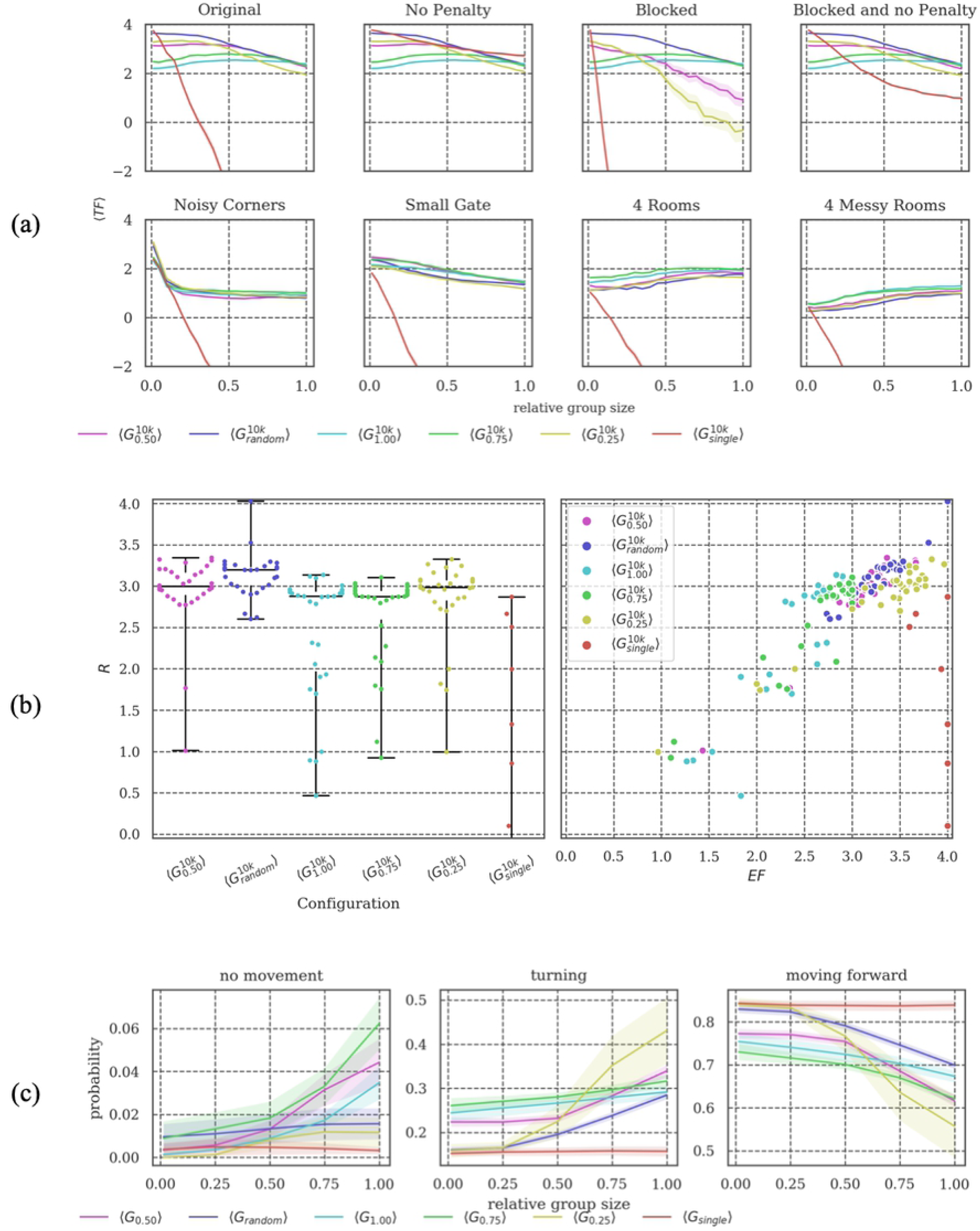
Reliability tests. **(a)** Overall, only ***G_single_*** fails to generalize across group sizes, as animats evolved without other group members did not develop strategies to avoid collisions (compare *Original* to *No penalty* test condition, where ***G_single_*** performs well throughout). There is a large difference in the *Blocked* environment between ***G_random_***, ***G_0.25_***, and ***G_0.50_***, while in other environments their task fitness is comparable, pointing to somewhat different navigation strategies. **(b)** On average, ***G_random_*** is the most reliable condition, followed by ***G_0.50_*** and ***G_0.25_***. Except for ***G_single_***, ***EF*** correlates with ***R*** in all groups. **(c)** Note that ***G_0.50_*** and ***G_0.25_*** change their behavior more with increasing animat density compared to ***G_random_***.

**Table 2:** Overview of the eight environments in which reliability tests were performed. They differ in environmental conditions and in the complexity of the world design.

| Label | Environmental Conditions | Environment (see Methods) |
| --- | --- | --- |
| Original | Active penalty<sup>1</sup>, no blocking<sup>2</sup> | See Fig 16(a) |
| No Penalty | **No penalty, no blocking** | See Fig 16(a) |
| Blocked | **Active penalty, active blocking** |  |
| Blocked and no Penalty | **No penalty, active blocking** |  |
| Noisy Corners<br>Small Gates<br>4 Rooms<br>4 Messy Rooms | Active penalty, no blocking | See Fig 16(b)<br>See Fig 16(c)<br>See Fig 16(d)<br>See Fig 16(e) |

<sup>1</sup> If penalty is active, animats receive penalty when colliding into each other.
<sup>2</sup> If blocking is active, an animat cannot move onto the location of another animat.

Next, we quantified the reliability across group sizes as the average task fitness ***R = 〈TF〉_GS_*** in the “Original” test condition (in this case, the average is calculated across group sizes not simulations as indicated by the subscript “***GS***”, which stands for group size). Panel (b) shows the distribution of these reliability values (***R***) and their dependency on evolved fitness (***EF***). Panel (c) shows how the animats’ behavior depends on the relative group size, evaluating the probability of an animat to stand still (“no movement”), turn, or move forward in the “original” test environment.

**Complexity analysis:** Fig 4, Fig 7, Fig 10, and Fig 13 show two types of metrics for MB complexity: (a) the distribution of *integrated information* (***Φ^Max^***) [18, 19], and (b) the corresponding *number of concepts* (#***Concepts***(***Φ^Max^***)) [18] per condition. While there may be simpler, less computationally demanding options for evaluating the causal complexity of the evolved MBs (see [15,16,21]), the chosen measures are fairly well established [15,19,22] and are theoretically motivated as part of the formal framework of integrated information theory (IIT) [18]. Briefly, a “concept” in IIT is a system subset that has a causal role within the system—an intrinsic mechanism. A concept causally constraints both the past and future states of the system, and is irreducible to its parts. The number of concepts (#***Concepts***(***Φ^Max^***)) thus captures the number of internal functions performed by individual system elements and combinations of elements. ***Φ^Max^*** quantifies how much of the information specified by all the concepts in a set of elements would be lost under a partition of the system, and it will be high if the set of elements has many concepts (functional differentiation) that are also highly integrated. Both measures are evaluated for the most integrated system subset, thus the ‘max’ superscript. For details please refer to the original publication [18] and to [19] for an application of these measures to evolve MBs.

**Fig 4.**
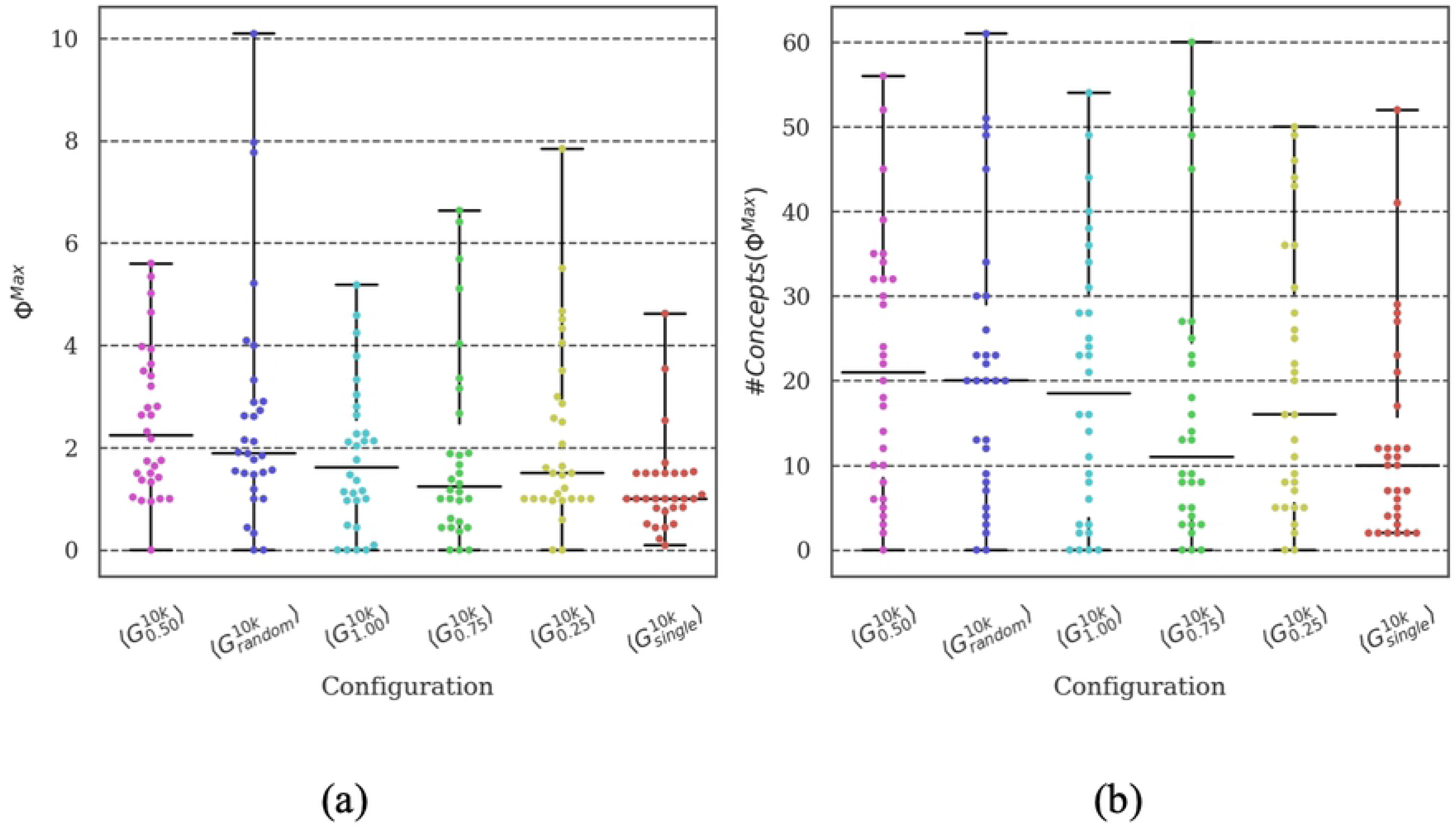
Distribution of brain complexity measures. Differences in **(a) *Φ^Max^*** and **(b)** the corresponding number of concepts was found between the most (***G_random_*** and ***G_0.50_***) and the least (***G_single_***) reliable setups. Due to the large variance in the data and the low sample size (30 simulations per evolutionary setup), differences in the mean between the remaining conditions did not reach statistical significance (see Supporting Information S3).

### Varying group size: Evolution under specialized conditions can produce reliable agents

In a first set of experiments, we compared animats that evolved within groups of fixed sizes (1-72 animats) (using the original animat and environment design in all cases). Preliminary results, including a comparison of the reliability of evolution conditions ***G_1.0-single_***, were presented in [17]. As shown in Fig 2(a) and reported in [17], group size during evolution does impact the animats’ ability to perform the gate crossing task (“task difficulty”) (Fig 1(a)), and it influences the final evolved fitness.

In our spatial-navigation task, single animats (group size of 1) frequently find an optimal solution within 10,000 generations, since colliding is impossible and walls (static obstacles) may guide the animat towards the gate. Increasing the number of animats in the environment makes it more difficult to navigate due to the penalty imposed upon colliding with another agent [17]. In our study, an animat was reliable if it could achieve high fitness under various conditions which they did not face during evolution. Reliability across group sizes was found to be high if the animats evolved in an environment where the density of animats was balanced (***G_0.5_*** and ***G_0.25_***) (see Fig 3(a,b) and [17]).

In our study, we included an additional comparison setup (***G_random_***), for which group size varied randomly during evolution, in order to explicitly evolve animats with high reliability. As shown in Fig 2(b), the final fitness values for ***G_random_*** were comparable to those evolution setups with fixed, intermediate group sizes (***G_0.5_*** and ***G_0.25_***) (though still significantly different (***p<.05)***, see Supporting Information S3 for all statistical tests).

It is no surprise that ***G_random_*** is the most reliable setup across varying group sizes (see Fig 3), since these animats already evolved under the conditions tested in the reliability evaluations. Notably, however, animats that evolved under specialized conditions with intermediate group sizes (***G_0.5_*** and ***G_0.25_***) are comparable to animats specifically evolved for reliability (***G_random_***) during evolution (see Fig 3). It is necessary to review the reliability tests in detail to observe differences between those evolutionary setups. ***G_0.50_*** and ***G_random_*** show similar reliability values ***R*** in the original environment setting, particularly for larger group sizes (> 50%) (see Fig 3(a)). Nevertheless, ***G_random_*** animats perform better with smaller group sizes, leading to comparable but still significantly different average ***R*** values (***p<.05***).

Of all test conditions (see Table 2), *Blocked* (in which animats cannot overlap) suggests a further difference between ***G_0.50_***, ***G_0.25_***, and ***G_random_*** (see Fig 3(a)): ***G_0.50_*** and ***G_0.25_*** are more severely affected by this deviation from standard settings in which animats can overlap, albeit under a penalty. While animats evolved in ***G_random_*** also experienced large group sizes with a higher likelihood of a penalty during evolution, ***G_0.50_*** and ***G_0.25_*** animats consistently faced only intermediate probabilities of colliding with other animats, which may have led to less effective strategies for avoiding collisions.

In addition to varying group sizes, we also tested the final generation of animats in four environments with different wall arrangements (Fig 3(a), bottom row). Performance decreased to similarly low levels in all conditions, but least for evolutionary setups with larger group sizes.

In terms of their behavior (Fig 3(c)), animats in ***G_random_*** were less idle and showed fewer turns and more steps forward in comparison with animats in ***G_0.50_***, particularly for large group sizes. This suggests that the behavior in ***G_random_*** is more fluid overall. By contrast, the specialized animats have to be more reactive to stay reliable, displaying larger difference in behavior across group sizes (see Table 3 for a more detailed explanation of the difference in behavior). Please refer to [17] for a more detailed discussion of behavioral differences across evolutionary setups with fixed group sizes ***G_1.0-single_***.

**Table 3:** Absolute difference between the state transition probability of ***G_0.50_*** and ***G_random_***. The first digit describes whether anything (wall or other animat) is sensed (1) or not sensed (0), and the second digit describes whether the animat moved/turned (1) or did not move/turn (0). Most notably, ***G_random_*** animats performed more movements even in the absence of sensor inputs than ***G_0.50_*** (“01→01”).

| $t \backslash SM$ | 00 | 01 | 10 | 11 |
| --- | --- | --- | --- | --- |
| 00 | 0.0000 | -0.0074 | 0.0000 | -0.0001 |
| 01 | -0.0079 | -0.0606<sup>1</sup> | 0.0136 | 0.0088 |
| 10 | 0.0005 | 0.0100 | 0.0063 | 0.0063 |
| 11 | -0.0001 | 0.0119 | 0.0031 | 0.0157 |

<sup>1</sup> Minus values indicate that the transition is more frequent in $G_{random}$ , while positive values indicate the opposite.

Fig 4 shows the distribution of ***Φ^Max^*** and ***#Concepts(Φ^Max^)*** [18, 19] as a measure of the complexity of the evolved MBs across evolutionary setups with different group sizes. While the most reliable evolutionary setups (***G_random_*** and ***G_0.50_***) do show the highest average values of ***Φ^Max^*** and the largest number of concepts (internal mechanisms), differences between conditions generally do not reach statistical significance (***p>.05***) due to the large variance in the complexity values (see Supporting Information S3). It would require more data (simulation experiments per evolutionary setup) to refine the mean of the intervals enough to verify the observed trend. In our predecessor study [17], a correlation of high reliability and task performance with high brain complexity was found using a simplified measure of brain complexity based on anatomical connectivity only. In addition, the integrated information measures employed here are sensitive to the causal interactions within the MBs. In the present data, significant pair-wise differences could be found between ***G_single_*** and the most reliable setups (***G_random_*** and ***G_0.50_***). As explained above, the task environment experienced by animats in ***G_single_*** is less demanding than for setups with larger group sizes. Our findings are thus in line with [19], which demonstrated higher ***Φ^Max^*** and ***#Concepts(Φ^Max^)*** for animats evolved in more complex environments.

### Varying cognitive design: Brain size and memory dependencies

In a second set of experiments, we used the same evolutionary setup as for ***G_0.50_*** in all tested conditions, but varied the number of available computational units in the animats’ MBs. In the baseline design ***G_0.50_***, it is possible to integrate motor units as memory units (by feedback loops to the hidden units, see Methods section). This was disabled in one condition ***G_!feedback_*** and therefore reduced the absolute capacity for memory from six to four binary units. Moreover, we designed animats with similarly small memory capacity but with feedback motors as a reference group (***G_smallbrain_***). Those animats had only two hidden units instead of four but the original type of motors with the possibility of evolving feedback loops. Again, the possible integration of motor units allows one to utilize information about past movements directly (*e.g., like the sensation of one’s legs*). Finally, we included a condition with larger MBs with eight hidden units and motor feedback (***G_bigbrain_***).

We observed that fitness and reliability across group sizes in the original environment decreased for animats with fewer computational units (see Fig 5 and Fig 6). However, while animats in ***G_smallbrain_*** still evolved to reasonably high fitness and reliability, ***G_!feedback_*** was lacking in both. This indicates that motor feedback facilitates evolution in our task environment. One behavioral difference between these two conditions was the reduced movement in the animats of ***G_smallbrain_*** (see Fig 6(c)). Furthermore, the state transition analysis shows that the motor units of animats in ***G_smallbrain_*** tend to change their behavior more often, while animats in ***G_!feedback_*** stay in the same state more often (see Table 4). Notably, ***G_!feedback_*** and, particularly, ***G_smallbrain_*** performed better than ***G_0.50_*** given large changes in the wall arrangement.

**Fig 5.**
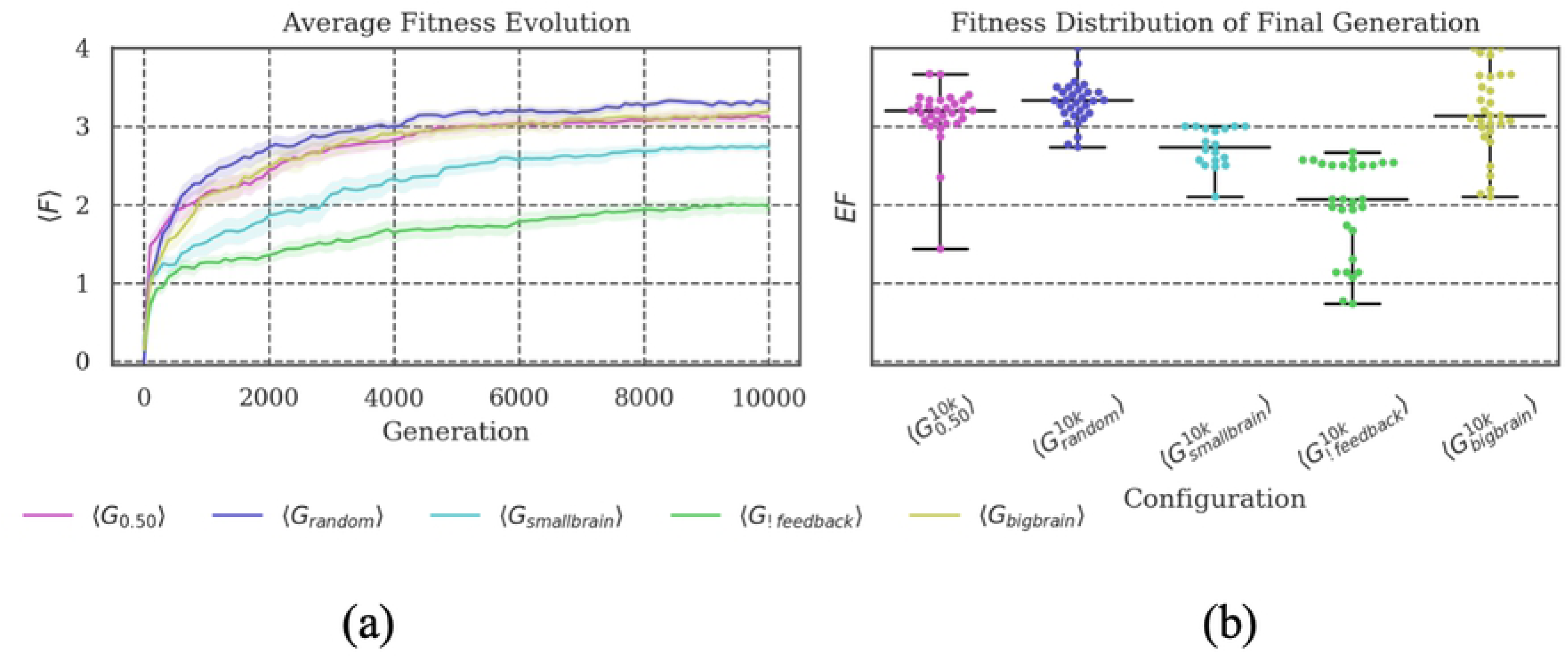
Fitness evolution and distribution of the final evolved fitness. **(a)** Less capacity for memory and internal computations impairs fitness evolution. Despite their similar capacity for memory, ***G_smallbrain_*** evolved higher fitness than ***G_!feedback_***. **(b)** Ceiling outlier suggest that animats in ***G_!feedback_*** are generally capable of performing as well as average animats in ***G_smallbrain_*** but that this is less likely. The performance of ***G_bigbrain_*** is comparable to ***G_0.50_*** with more distributed outcomes.

**Fig 6.**
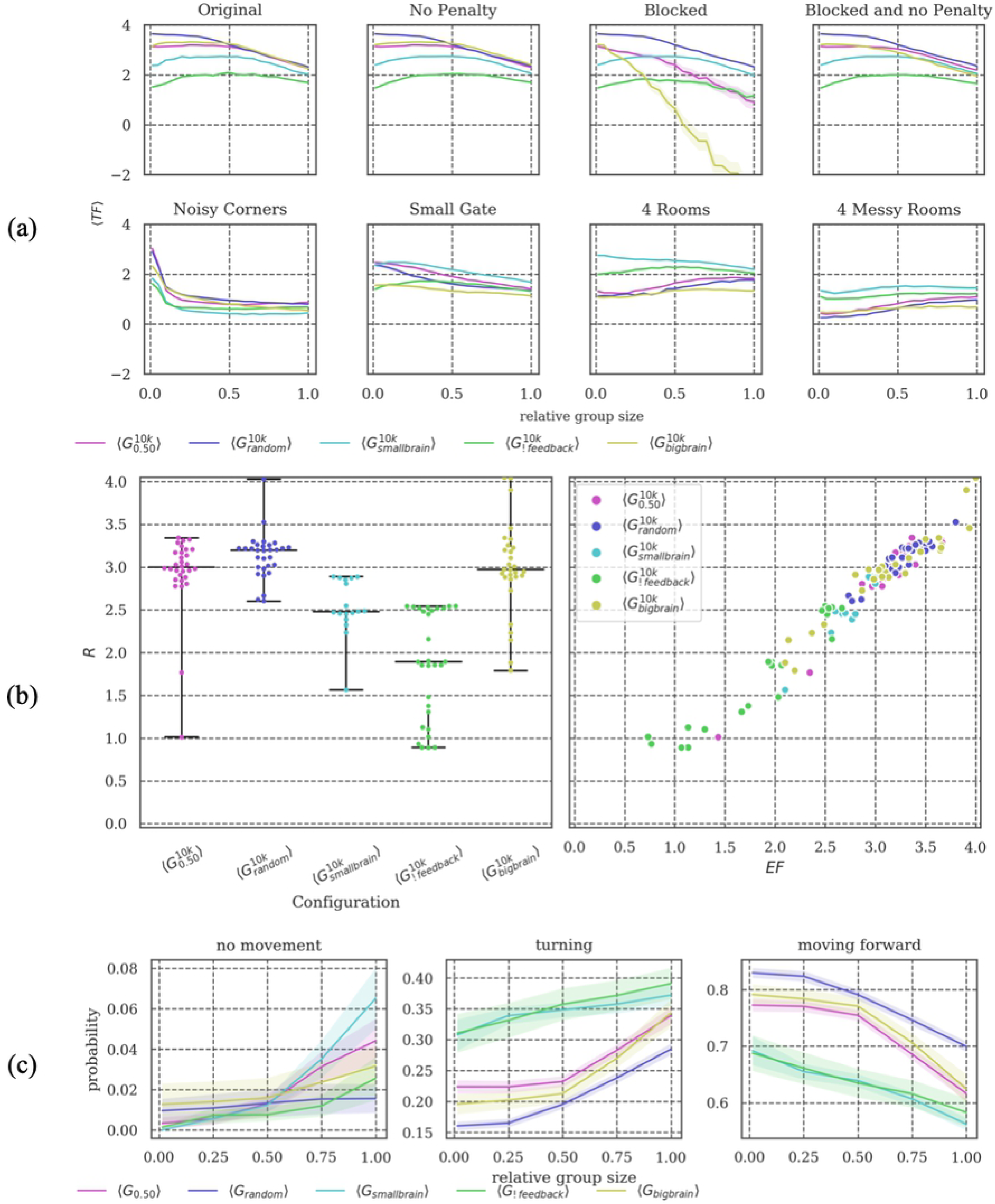
Reliability tests. **(a) *G_smallbrain_*** is more reliable than ***G_!feedback_***. Considering ***G_bigbrain_***, animats in this group are overall comparable to the baseline condition ***G_0.50_***, but show worse performance in the *Blocked* test condition and some of the modified environments for larger group sizes. **(b)** Reliability ***R*** correlates with ***EF*** for all setups. The lower reliability of ***G_smallbrain_*** and ***G_!feedback_*** compared to baseline can thus be explained by their already lower evolved fitness values. Note, however, that ***G_smallbrain_*** and ***G_!feedback_*** perform better than ***G_0.50_*** across group sizes in the *4 (Messy) Rooms* test conditions (see **(a)**). **(c)** For larger group sizes, ***G_smallbrain_*** remains static more often than ***G_!feedback_***.

**Table 4:** Absolute difference between the state transition probability of ***G_smallbrain_*** and ***G_!feedback_***. The first digit describes whether anything (wall or other animat) is sensed (1) or not sensed (0) and the second digit describes whether the animat moved/turned (1) or did not move/turn (0). Most notably, animats in ***G_smallbrain_*** switched more often between sensing and moving than animats in ***G_!Feedback_***(“01→10”, “10→01”, but “11→11”).

| $t \backslash SM$ | 00 | 01 | 10 | 11 |
| --- | --- | --- | --- | --- |
| 00 | 0.0000 | 0.0001 | 0.0000 | 0.0000 |
| 01 | 0.0000 | -0.0167<sup>1</sup> | 0.0237 | -0.0046 |
| 10 | 0.0000 | 0.0194 | 0.0011 | 0.0029 |
| 11 | 0.0001 | -0.0004 | -0.0015 | -0.0241 |

<sup>1</sup> Minus values indicate that the transition is more frequent in $G_{feedback}$ , while positive values indicate the opposite.

By contrast, more hidden units (***G_bigbrain_***) do not improve average fitness or reliability in any of the tested conditions (see Fig 5 and Fig 6). While ***G_bigbrain_*** overall seems very similar to the baseline setup ***G_0.50_***, differences can be observed in the *Blocked* and *Small Gate* test conditions (see Fig 6(a)). In principle, more computational units allow for better performance. However, the larger space of possible solutions may also impede fitness evolution (note the larger variance for ***G_bigbrain_*** compared to ***G_0.50_*** in Fig 5(b) and Fig 6(b)).

Considering brain complexity, the evolutionary setups with smaller MBs (***G_smallbrain_*** and ***G_!feedback_***) have significantly lower ***Φ^Max^*** and fewer concepts than the baseline condition (***G_0.50_***). Between those two conditions, ***G_smallbrain_*** shows significantly higher ***Φ^Max^*** and more concepts as compared to ***G_!feedback_*** (see Fig 7). This correlates with the larger evolved fitness values of ***G_smallbrain_*** in Fig 5 and its associated higher reliability in Fig 6. Note that calculating ***Φ^Max^*** and the corresponding number of concepts was not possible for ***G_bigbrain_*** since exhaustive evaluations across many systems and states are not currently feasible when using the *pyphi* software package to compute measures of integrated information theory for networks of that size (>10 units) [23].

**Fig 7.**
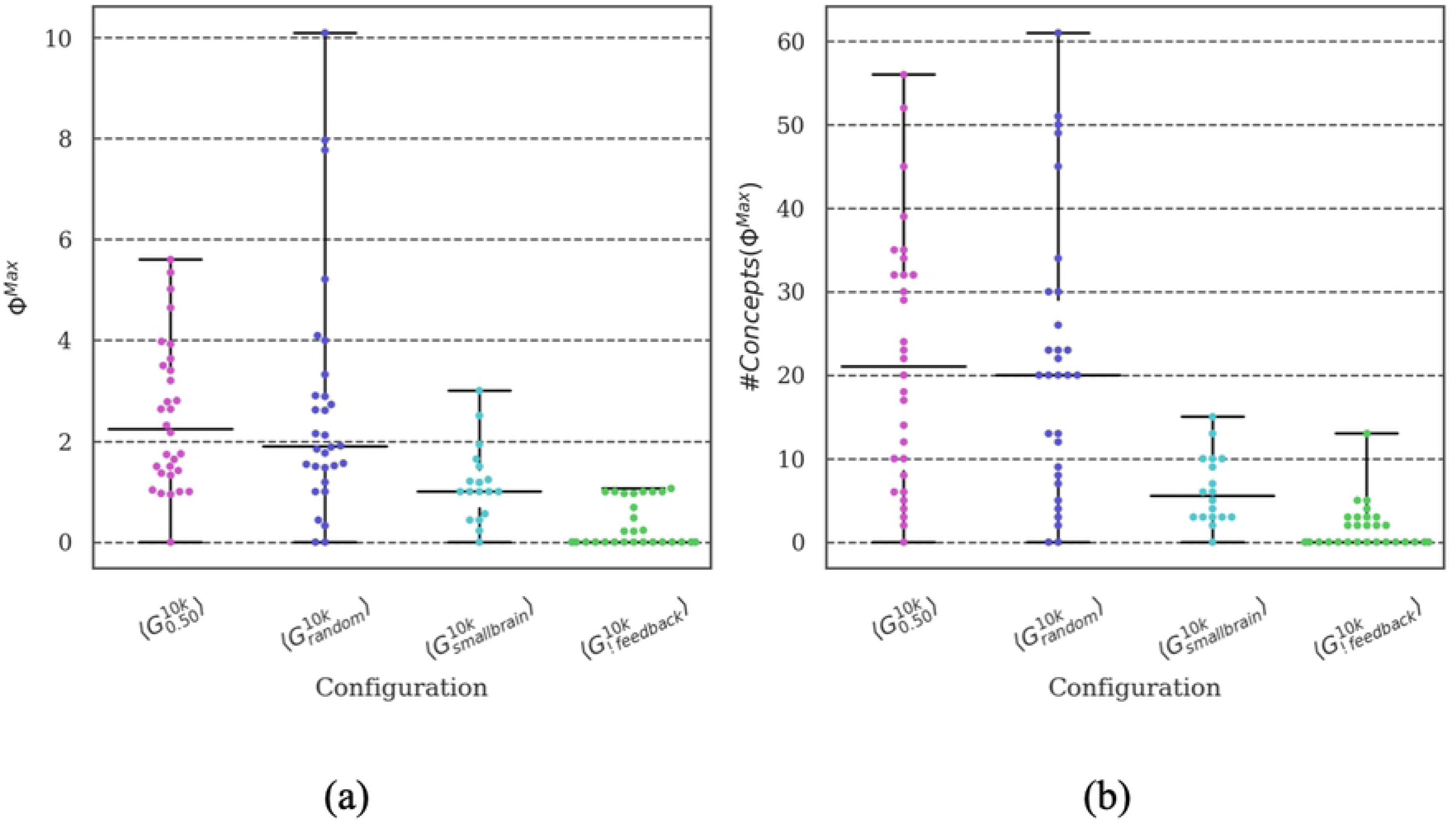
Distribution of brain complexity measures. Compared to the baseline, the smaller MBs (***G_smallbrain_*** and ***G_!feedback_***) have lower ***Φ^Max^*** and fewer corresponding concepts. Animats in ***G_smallbrain_*** show higher ***Φ^Max^*** and have more corresponding concepts compared to ***G_!feedback_*** animats, many of which have ***Φ^Max^ = 0***. Due to computational reasons, the brain complexity of ***G_bigbrain_*** could not be calculated (see text).

### Varying interaction conditions: Evolution of beneficial interaction

In our evolution simulations, the fitness function used for selection depended on the average task fitness of all animats in the group. Moreover, individuals received penalties for colliding with other group members. Since it is hardly possible to directly observe cooperative interactions, we used a third set of simulations to manipulate aspects of the fitness function and physical interaction between animats to identify to what extent these features influence both the fitness and the reliability. For this purpose, we considered four different evolutionary setups besides the baseline setup ***G_0.50_***: ***G_single_*** (same as above), ***G_!penalty_***, ***G_blocked_***, and ***G_blocked/!penalty_*** (see Table 1 for a detailed description). ***G_random_*** is also included in the figures for comparison.

Among the novel setups, only animats in ***G_blocked_*** were subject to the collision penalty during evolution, whereas later, during the *Original* reliability tests, all conditions were subject to a penalty. Not being able to share the same position (as in ***G_blocked_***) hardly influenced the final fitness, reliability, or behavior of the evolved animats (Fig 8 and Fig 9, compared to the baseline condition). ***G_!penalty_***, where reacting to other animats had no direct effect on the fitness evolution, showed very similar fitness evolution, reliability curves, and behavior to ***G_single_***. Considering the reliability tests in Fig 9(a), the top row shows the reliability across group sizes in the *Original* environment, and under varying interaction conditions: *No Penalty*, *Blocked*, and both *Blocked and no Penalty* (from left to right). In the bottom row of Fig 9(a), animats are evaluated under the same interaction rules as they evolved in while only facing a modified environment.

**Fig 8.**
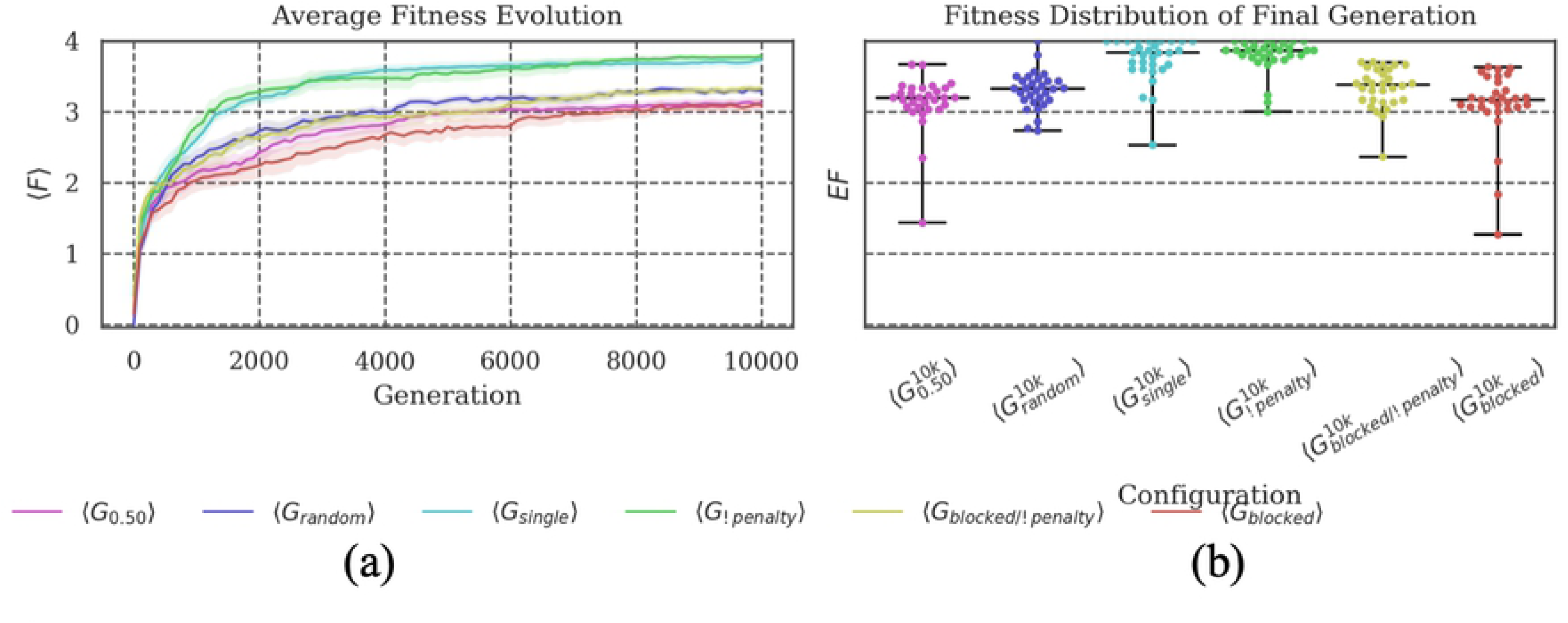
Fitness Evolution and distribution of the final evolved fitness. The animats in conditions without a penalty (***G_blocked/!penalty_*** and ***G_!penalty_***) evolved to relatively high fitness levels. In particular, ***G_!penalty_*** evolved like ***G_single_***, since animats in both conditions were not impacted at all by other animats. Similarly, ***G_blocked_*** seemed equivalent to the baseline setup ***G_0.50_***, while ***G_blocked/!penalty_*** evolved to slightly higher fitness values, comparable to ***G_random_***.

**Fig 9.**
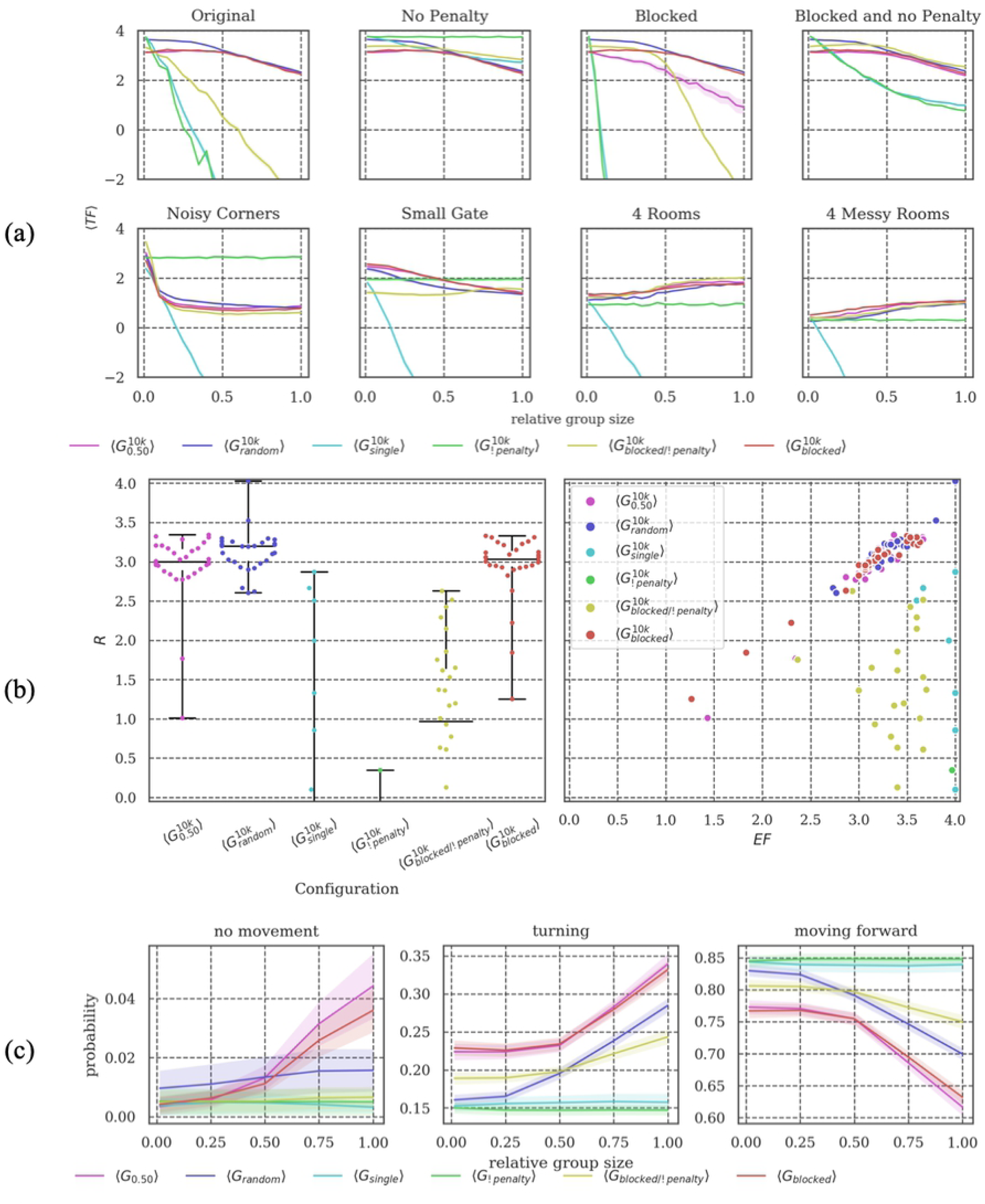
Reliability Tests. **(a)** There was a significant difference between animats in ***G_blocked/!penalty_*** and animats in ***G_!penalty_***. Being blocked was essential for retaining some reliability if no penalty was given. **(b) *G_!penalty_*** showed similar reliability as ***G_single_***, whereas ***G_blocked_*** showed similar reliability as ***G_0.50_***. **(c)** These similarities were also reflected in the animats’ behavior. The behavior of animats in ***G_blocked/!penalty_*** was more reactive to changing group size than ***G_!penalty_***.

Comparing the reliability tests of ***G_blocked/!penalty_***, ***G_blocked_*** and ***G_!penalty_*** (Fig 9), we observed significant differences between the setups, which let us assume that there is implicit cooperation. In this context, we want to highlight that ***G_!penalty_*** performs relatively poor for larger group sizes in the environment designs with large modifications (in *4 (Messy) Rooms*) as compared to the other setups. This is an indicator for the evolution of beneficial interactions between group members in evolutionary setups with a collision penalty and/or blocking. The decline in task fitness of ***G_blocked/!penalty_*** for higher group sizes under test conditions with a collision penalty showed that these animats did not avoid physical interactions with their group members, while ***G_blocked_*** animats were generally comparable to ***G_0.50._*** However, even ***G_blocked/!penalty_*** animats had an advantage compared to ***G_!penalty_*** in the *4 (Messy) Rooms* environment, which may be due to some implicit form of cooperative behavior.

Considering the brain complexity of animats in ***G_blocked_*** and ***G_blocked/!penalty_***, we can report similar values to ***G_0.50_***(see Fig 10). Whether animats received a penalty for crossing each other, or whether crossing was prohibited to start with, did not significantly affect their evolved fitness, reliability, behavior, or brain complexity. Likewise, the brain complexity measures for ***G_!penalty_*** were comparable to those of ***G_single_***, in line with the behavioral results above.

**Fig 10.**
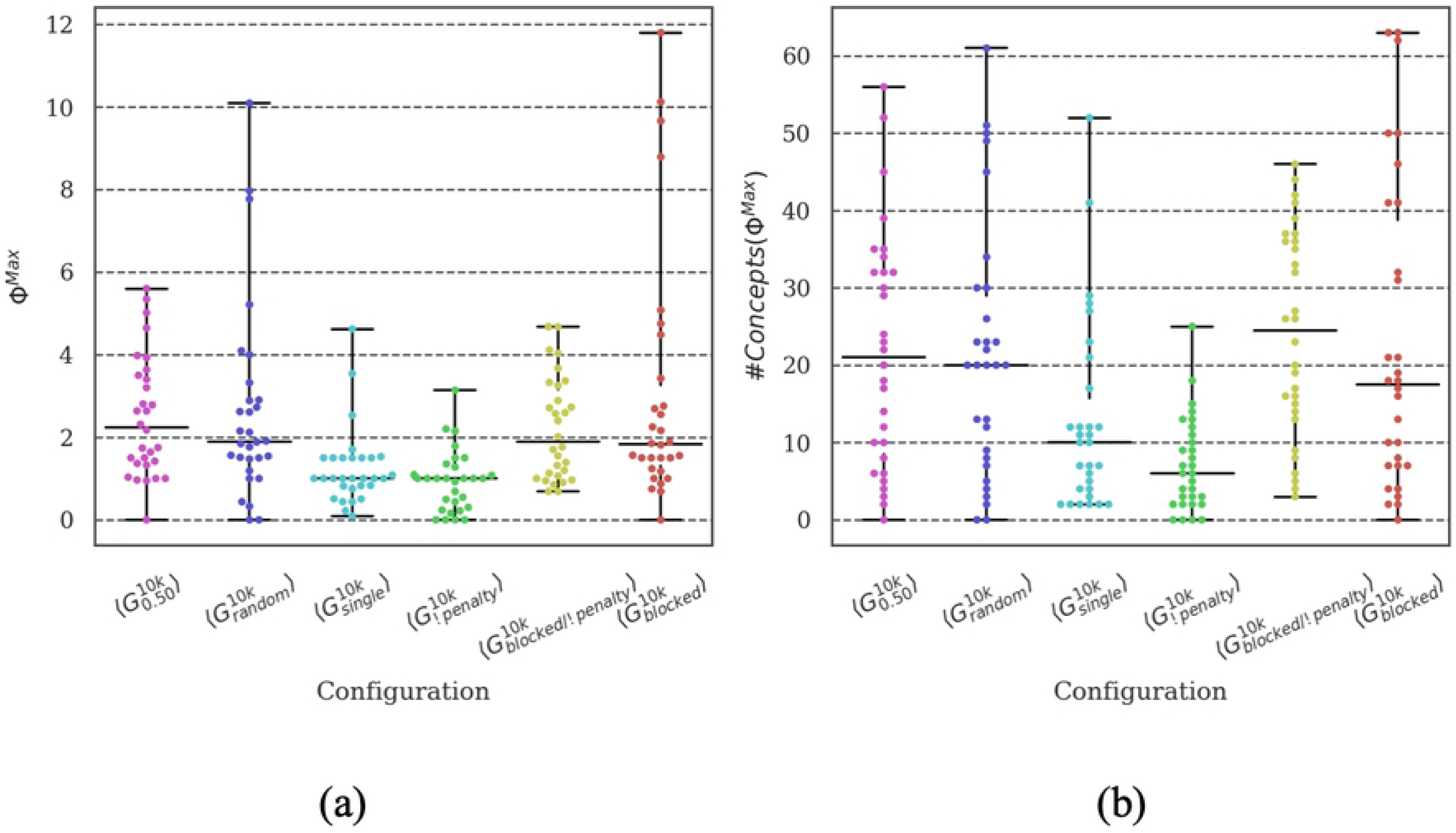
Distribution of brain complexity measures. In evolutionary setups where crossing each other was not possible (***G_blocked_*** and ***G_blocked/!penalty_***), the brain complexity was comparable to the complexity of ***G_0.50_***. By contrast, animats in setups where the reaction to fellow animats had no reasonable effect on their performance (***G_single_*** and ***G_!penalty_***) showed lower brain complexity. Still, there was high variance in the data of brain complexity.

### Varying sensor configuration: Sensory capacity influences reliability and intrinsic complexity

Finally, we manipulated the ability of dealing with the task (task difficulty) by changing the sensor configuration of the animats. In addition to the baseline architecture, we designed animats with sensors on three sides ***G_3sides_*** (front, left and right), without an agent sensor ***G_!agent_*** and with a universal sensor ***G_w=a_*** (sensing wall and agent as indiscriminate obstacles). Fig 11 reveals that it is necessary to have the ability to sense nearby animats, and be able to differentiate between walls and animats, in order to achieve reasonable fitness values. Generally, it was an advantage to be equipped with sensors on more sides for both high task fitness and high reliability.

**Fig 11.**
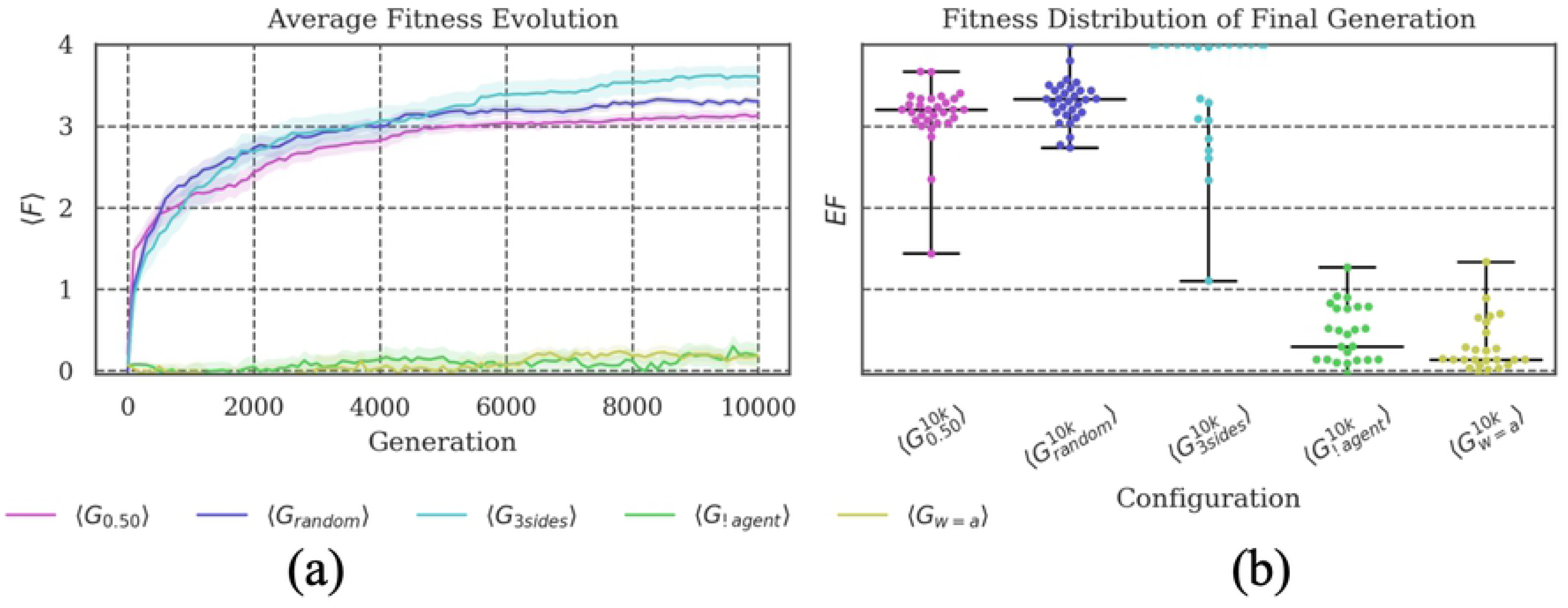
Fitness Evolution and distribution of the final evolved fitness. The average evolved fitness showed that animats in evolutionary setups without specific sensors for other animas (***G_!agent_*** and ***G_w=a_***) achieved no reasonable fitness. By contrast, animats in ***G_3sides_*** outperformed ***G_0.50_***, and ***G_random_***, but also had more outliers with lower fitness and performed worse than the baseline condition in early generations (up to ∼10k generations).

Regarding reliability, we would first like to highlight animats in the ***G_3sides_*** condition. They consistently outperformed the animats in other groups except in two test conditions: *Blocked* and *Noisy Corners* (see Fig 12). This shows that animats which are equipped with more sensors do have an advantage on average, but they may also perform worse than animats with fewer sensors under some circumstances. The sensory signals in these specific environments might have been too different from the information patterns the animats evolved in and were thus specialized for.

**Fig 12.**
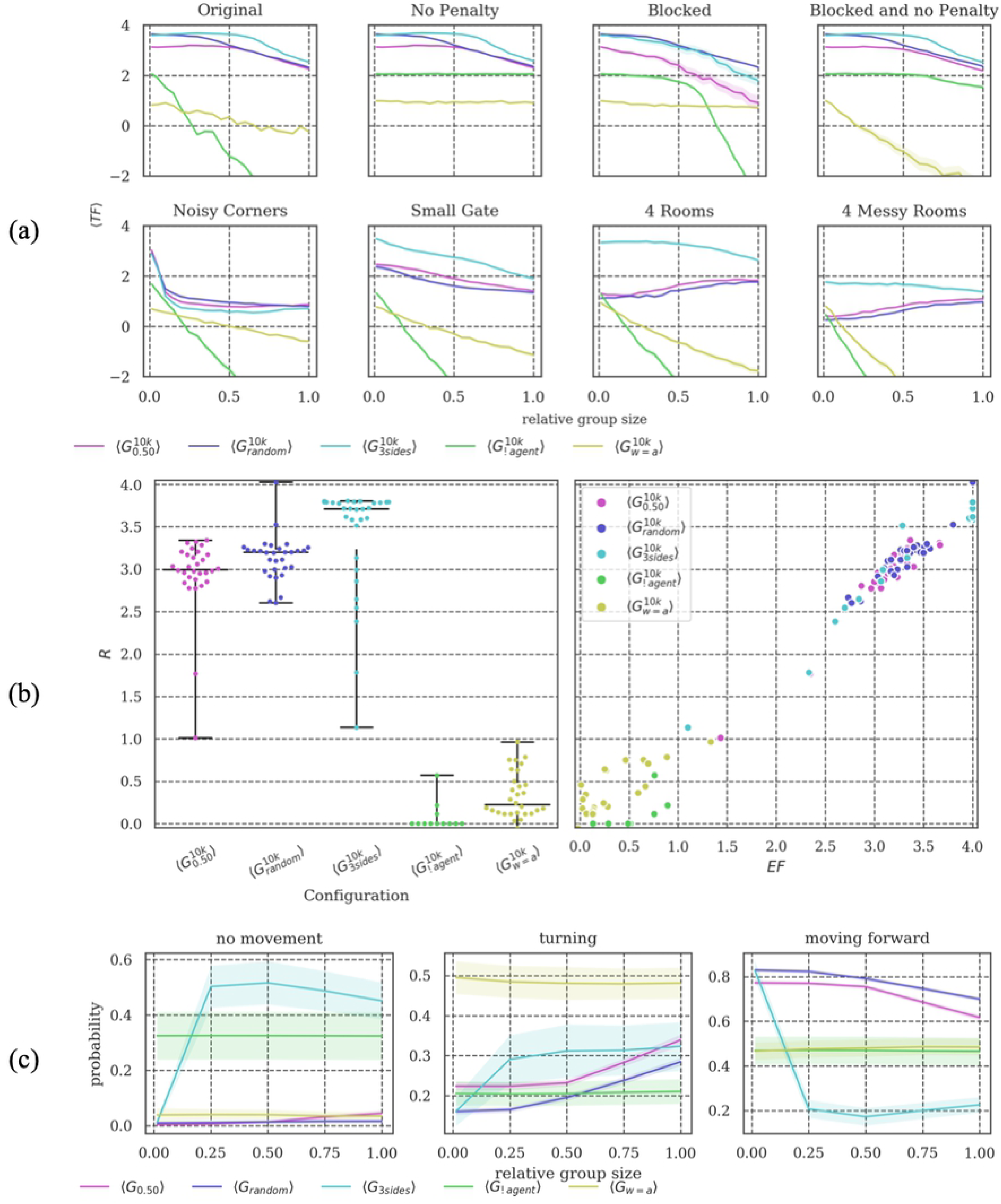
Reliability Tests. **(a-b)** The ***G_3sides_*** condition was the most reliable in most test conditions, except in *Blocked* and *Noisy Corners.* In terms of reliability, sensing everything (***G_w=a_***) with one sensor is still better than only sensing the walls due to a missing animat sensor (***G_!agent_***). **(c)** Setups with few sensors evolved no general behavior (high variance of movement between the 30 different evolutions, shaded area). The ***G_3sides_*** setup becomes more reactive as soon as the animat density starts to rise.

Opposite behaviors can be observed for the animats in ***G_w=a_*** and ***G_!agent_***. In this case, animats were not evolving to reasonable fitness values. Nevertheless, we could observe differences between the two conditions from their reliability values. While ***G_w=a_*** animats had only one sensor which does not discriminate between the wall and other animats, ***G_!agent_*** was missing the animat sensor completely. ***G_!agent_*** showed better task fitness than ***G_w=a_*** in test conditions with small group sizes and without a penalty. Considering the evolved behavior, ***G_w=a_*** animats (Fig 12(c)) were not reactive to other animats, which suggests that they did not evolve the capacity to differentiate between the animats and the walls internally, e.g., through memory.

Analyzing the brain complexity showed that animats equipped with fewer, but also with more sensors than in the baseline setup ***G_0.50_*** evolved MBs with lower complexity (see Fig 13), albeit for different reasons. Based on the very low evolved fitness for ***G_w=a_*** and ***G_!agent_*** (see Fig 11) we can conclude that their MBs did not develop the necessary structure and mechanisms to solve the task, as reflected by their low brain complexity. By contrast, animats in ***G_3sides_*** achieved high performance and reliability, but did not evolve any integrated information (***Φ^Max^ = 0***) in many cases. This observation was in line with previous findings on the relation between sensory capacity and internal complexity [19] and suggested that high brain complexity in cognitive systems depends on a need for internal memory and computation, which may decrease if an animat is equipped with more sensors. Please refer to the next section for a general discussion about the relationship between task performance, reliability, and brain complexity.

**Fig 13.**
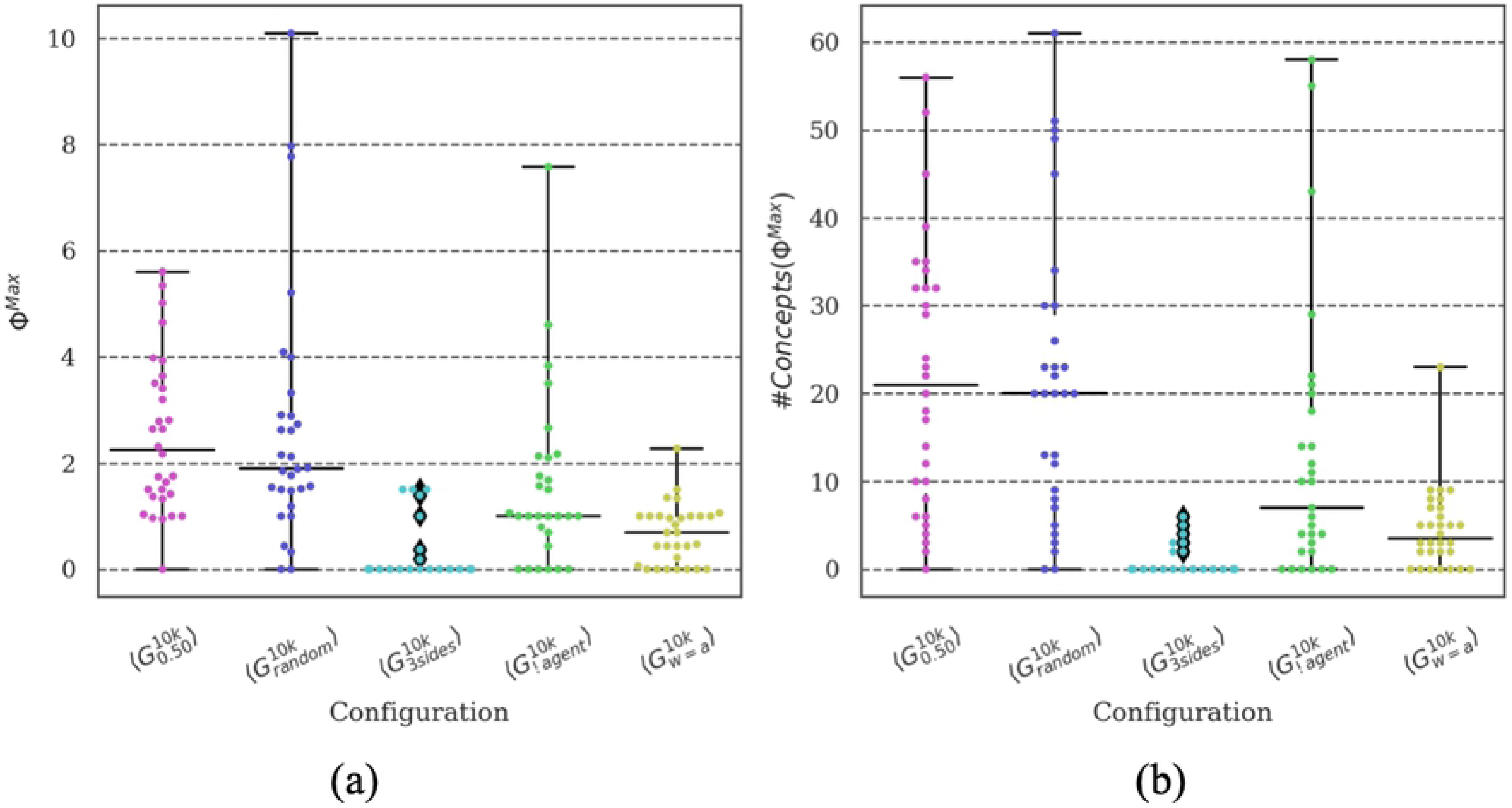
Distribution of brain complexity measures. Animats in the ***G_3sides_*** condition showed the lowest brain complexity of all setups despite having the highest evolved fitness and reliability. By contrast, animats with limited sensor information (***G_!agent_*** and ***G_w=a_***) had lower than baseline complexity values, but also low evolved fitness (***EF***, see Fig 11).

## Discussion

The evolution of cooperative multi-agent systems might be the next frontier in the context of evolving artificial agents, in which context not much is yet known about conditions that give rise to cooperative behavior and the complex inter-dependencies between individual and group goals [24]. For example, there might be many factors that influence whether the individuals either bow to the group or act by egoistic rules [25]. In this study, we used animats equipped with MBs (introduced by Edlund et al. [21]) to study how group performance and its reliability under modified conditions depended on the individual, interactions between individuals, as well as specific features of the MBs’ evolution.

### Prior work investigating group evolution

Earlier research that implemented groups of MBs concentrated on predator-prey environments and showed that animats can (co-)evolve swarm behaviors [26–28]. The animat design in this work was generally based on a design in Marstaller et al. [15], who evolved individual MBs with the goal of solving perceptual-categorization tasks. Another method of simulating swarm behavior is neuro-evolution, i.e., the evolution of *artificial neural networks (ANN)* [29–31]. As in Olson et al. [27], these neuro-evolution experiments produced agents which evolve in a swarm to solve a predator-prey task.

Other researchers have investigated the effect of group size in the evolution of groups of simulated agents beyond predator-prey scenarios in a more general context. They find that the behavior of the group of agents and the individual agent is dependent on the group size [32, 33]. In another study which changed the group size during evolution, the authors show that it can be easier for smaller groups than larger ones to organize themselves [5].

The effect of changing swarm sizes has also been investigated in the context of natural biological systems: Brown [25] examined which factors are decisive for the individual to either join a swarm or behave egoistically. The study focused on experimenting with environmental qualities and swarm size. Brown defined *optimal swarm size* as the best trade-off between the advantage of balancing costs between individuals in the swarm and the disadvantage of sharing the resources (energy/food) with the whole swarm. In an earlier study, Pacala et al. [4] report that swarm size constrains information transfer and task allocation. They argue that the information exchange varies and the task allocation changes, depending on the swarm size of ant-colonies. Pacala et al. [4] also argue that swarm behavior is the product of social interaction, individual interaction, and the interaction with the given environment. In a more recent work [34], we found arguments that swarm behavior arises if there is sufficient density within the swarm.

### Factors that impact task performance and reliability

In line with the variety of dependencies identified in these earlier studies, our simulation results suggest that group performance and reliability under modified conditions are complex multidimensional phenomena. Our work is illustrative, as it shows that there is high complexity even in the simplified experimental setting of small artificial organisms evolving within a particular evolutionary setup which is completely controlled by the experimenter. Nevertheless, by creating a variety of environments and animats, we were able to identify several factors that influence fitness evolution and post-evolutionary reliability.

Generally, task difficulty (the ability to evolve high fitness in a given task environment) depends on the complexity of the environment, but also on the animats’ architecture (see also [19]). In the specific evolutionary setup investigated here, evolved fitness negatively correlated with group size as a result of the imposed penalty for collisions (see Fig 2). On the other hand, animats evolved in fixed, intermediate group sizes are most reliable to changes in group size, and, in fact, comparable to animats evolved for reliability that experienced random group sizes during evolution (Fig 3(b)). Yet animats evolved in large groups performed slightly better in modified environments (Fig 3(a), bottom row). A similar trade-off can be observed for different animat architectures: animats with less capacity for memory (***G_smallbrain_*** and ***G_!feedback_***) evolved to lower fitness levels than the baseline condition (***G_0.50_***) (Fig 5), and were less reliable under changes in group sizes but still showed better performance in some of the modified environments (Fig 6(a)). More hidden units (***G_bigbrain_***) did not provide further advantages compared to ***G_0.50_***. Finally, more sensors (***G_3sides_***) proved advantageous for both evolved fitness and reliability under almost all modified test conditions. However, even ***G_3sides_*** performed worse than the baseline in one of the modified environments (*Noisy Corners*). Within most specific environmental setups, reliability to changes in group size was, moreover, correlated with evolved fitness (Figs 3/6/9/12 (b), right panel).

Overall, we found that the right balance is essential: If the environmental design is balanced to the animats’ architecture (having the right sensor setup, memory capacity, and motor setup), animats evolved consistent reliability, even if it was not specifically trained for. In other words, animats that were well-equipped for dealing with their original task environment (and thus achieved high evolved fitness) were generally also able to remain reliable given small modifications to task conditions. However, evolutionary setups that seem less adapted (lower evolved fitness) overall may still have advantages under some conditions.

### Interactions between individuals in the group

In this study, we did not explicitly implement any form of direct communication between animats. Nevertheless, through triangulation, we can partly answer whether the evolutionary setup we employed here may have led to the evolution of implicit cooperation between group members. To that end, we have shown that it was necessary for animats to perceive their fellow group members, and that they use this information to achieve reasonable evolved fitness and reliability (Fig 11 and Fig 12). Moreover, animats evolved in large groups showed an advantage across group sizes in modified environments (Fig 3(a), bottom), while animats that evolved without a collision penalty (***G_!penalty_***) performed worse in some of the modified environments, even if tested without a penalty (Fig 9(a), *4 (Messy) Rooms*).

Hypothetically, this type of implicit interaction between animats is less related to *verbal communication*, but it may relate more to communication through behavior (e.g., like bees performing their dance). As we know from previous studies, swarm behavior in nature can also be the result of simple reactions to local neighbors [3, 35]. We argue that animats are interdependent in this way, even if there is no explicit information exchange between them. The observed instances of cooperative behavior can thus be viewed as an emergent phenomenon of the evolutionary process.

### Relation between brain complexity, task performance, and reliability

Previous studies applying measures of integrated information to adaptive animats equipped with MBs [19,21,36] have observed that ***Φ^Max^*** and related measures on average increase over the course of evolution, which correlates with increasing task performance (see Table S6 in Supporting Information S2). Moreover, as demonstrated in [19], this increase depends on the complexity of the task environment relative to the animats’ sensor capacity: MBs that evolved in task environments which required more memory and internal computation developed, on average, higher ***Φ^Max^*** values and a higher number of concepts.

For the evolutionary setups with the standard animat architecture as in ***G_0.50_***, we found the highest values of ***Φ^Max^*** and #***Concepts***(***Φ^Max^***) for medium group sizes ***G_0.50_***, and ***G_blocked_***, and for ***G_random_***. These setups were also among the most reliable across group sizes (see also [17] for similar results using a simplified measure of brain complexity). By contrast, significantly lower ***Φ^Max^*** values were found for ***G_single_*** and ***G_!penalty_***, the two setups in which task fitness during evolution did not depend on interactions with other animats. As argued above, ***G_single_*** and ***G_!penalty_*** thus effectively evolved within a simpler task environment than ***G_0.50_***, ***G_blocked_***, and ***G_random_***, which explains their lower ***Φ^Max^***.

Compared to ***G_0.50_***, evolutionary setups with altered animat architectures showed consistently lower values of ***Φ^Max^*** and #***Concepts***(***Φ^Max^***). Limiting the animats’ sensor capacity (***G_!agent_*** and ***G_w=a_***) or the number of available memory units (***G_smallbrain_*** and ***G_!feedback_***) interfered with their capacity for successful evolution in the spatial navigation task. Their lower performance was thus accompanied by less developed MBs with lower ***Φ^Max^*** and fewer concepts. Given more time to evolve (more generations), both their performance and their brain complexity might still increase. By contrast, more sensors allowed for better performance based on high amounts of external information, which effectively decreased the need for internal complexity (memory and computations) and thus may also lead to low ***Φ^Max^***, as observed here for ***G_3sides_***.

In theory, high fitness in any given environment could be achieved without information integration (e.g., by a system with a large feed-forward architecture [18]), and information integration can be high even if there is no reasonable fitness, which partially explains the large variance in the brain complexity measures (see, e.g., outliers for ***G_!agent_*** in Fig 13) However, given a certain requirement for memory and context sensitivity, constraints in the number of sensors and hidden elements may give rise to an empirical lower boundary on the amount of integrated information necessary to perform a given task [19,21,36,37].

In summary, for a given MB architecture, higher brain complexity seems to be related to better performance and reliability. However, future work should explore under which environmental conditions additional sensors, or more internal units, become more advantageous for the evolution of higher task performance and reliability.

### Limitations

Our work modeled one particular, small-scale scenario. Future work should consider other task environments which may strengthen the generality of our results. Moreover, further evolution or training scenarios for artificial organisms should be considered as well.

While the measures that we employed to assess the complexity of the evolved MBs are theoretically motivated [18], they are also computationally very complex. This made it difficult to evaluate a larger number of evolution simulations in order to achieve better statistical power. This is why alternative, approximate measures should be considered, too. For instance, the *largest strongly connected component* (and other graph metrics) can be used as a proxy for system integration and thus brain complexity [17]. Efficient approximations would also enable investigation into how brain complexity develops across generations. Moreover, ***Φ^Max^***, and the associated number of concepts, are causal measures that assess the degree to which the mechanisms within a MB are differentiated and integrated. Future work should also consider and explore alternative informational or dynamical measures [e.g., 38– 40]. In this study, we concentrated on the reliability tests, so the brain complexity analysis was not the subject of more in-depth investigation.

### Conclusion

It is challenging to remain reliable in a dynamic and volatile world while also trying to succeed in a given task. So, investigating the characteristics of this reliability might help to develop implications and strategies for improving reliability. We showed that reliability is a complex concept to investigate, especially when considering not only individuals but an organized group. Yet we were able to isolate essential influencing factors to better understand the positive and negative effects of changing group size, environment design, and individual cognitive ability on task reliability. This research asserts that task efficiency and effectiveness is not the only goal; task reliability is also worth striving for. We have also offered a computational approach for investigating this concept.

## Materials and Methods

We used an EA to generate simulated animats evolving in groups, and defined and tested various animat architectures and evolutionary environments to evolve animats having heterogeneous behavior, fitness, and reliability. Afterwards, we conducted post-evolutionary tests to assess the reliability of the different evolutionary setups. This section explains the animat designs, the environment, the evolutionary simulations, and the experiment setup. We used *MABE (Modular Agent-Based Evolver)* [41] as a computational evolution framework with the same parameters as in previous work [17] (see Table S7 in Supporting Information).

As we state in the introduction, we studied the changes in behavior and task performance of evolved animats while manipulating environmental and cognitive conditions, which also changed the ability to achieve the goal of the task (task difficulty). The idea was that the individual animat had to solve a two-dimensional spatial-navigation task, thus forcing individuals to react to other animats in order to reach a high fitness value. This task was a redesign by Fischer et al. [17] of a task environment initially developed by Koenig et al. [20]. An animat can usually differentiate between static (borders and walls) and dynamic objects (animats) in the environment through two distinct sensors. This design allowed for the evolution of social behavior based on passive interactions between animats (we observed, e.g., “waiting”, or “following” behavior).

### Animats Architecture

The EA evolves animats with MBs, which contain a set of discrete, binary computational units (“neurons”). Each unit has its own update rules receiving inputs from and sending their output to other units. In this study, the decision system (the connectivity between units and their update-rules) was implemented by *Hidden Markov Gates (HMGs)*. The HMGs connect the nodes of the MB indirectly. Fig 14 visualizes a simple example, in which an HMG is connected to four units. The decision system inside an HMG can be diverse. In this research, we evolved discrete lookup tables. The lookup tables translate the states of the connected input units at ***t*** to the new states of connected output units at ***t+1***. The motor or memory units can represent the output units of the HMG. In this study, the EA evolved genomes with a string of natural numbers. The individual numbers encoded the HMGs: the number of HMGs, the lookup tables, the connected input units, and the connected output units. The EA mutated the genomes in each generation. Each locus in the genome mutated with a certain probability. In addition, larger sections could be deleted or added to the genome [21, 42] (again, all parameters are listed in Table S7 within the Supporting Information).

**Fig 14.**
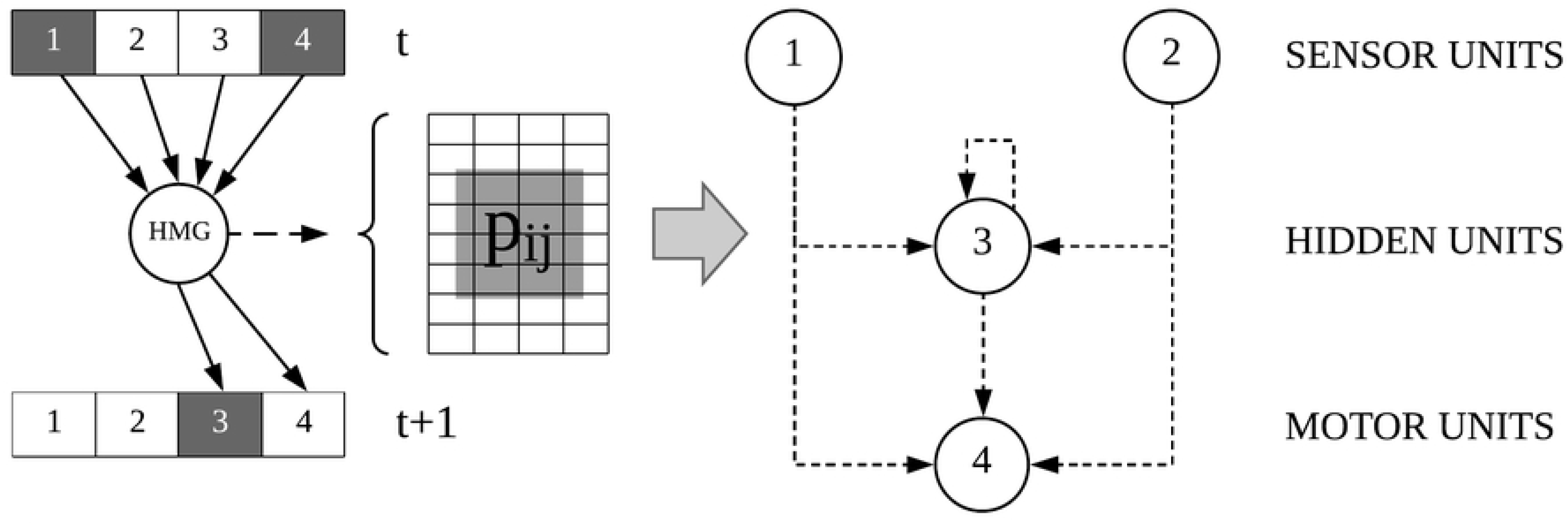
Example of an MB. An MB [21] has three components: (1) Units with a binary states (“1”- “4”), (2) HMGs and (3) the connections between the binary units and the HMGs. The connections between the units can be derived from the connections to the HMGs. HMGs contain the mechanism, e.g., a probabilistic lookup table, to transform the brain state of units at ***t*** to the state at ***t+1***.

All units in the animat’s MB have binary states, either ***1*** or ***0***, e.g., a sensor turns ***1*** if an obstacle is detected and a motor switches to ***1*** if it is active. Two motors provide the ability to turn 90 degrees left or right, and to move forward (if both motors are in state ***1***). Since the units within a MB can be interconnected in a recurrent manner, they have the potential to create internal memory. We evolved animats with five different animat designs. Fig 15 gives a schematic overview of all animat designs. In addition to the baseline cognitive architecture, which was introduced already in [17], further deviations were designed to investigate the influence of different cognitive setups on the resulting evolved behavior, task performance, and reliability. The sensors had a detection range of one unit. Typically, the motor units could also feedback to the hidden and motor units, thus acting as additional brain capacity, since knowledge about previous motor states is directly available for computing the next state. Additionally, we designed an animat without motor feedback (***G_!feedback_***).

**Fig 15.**
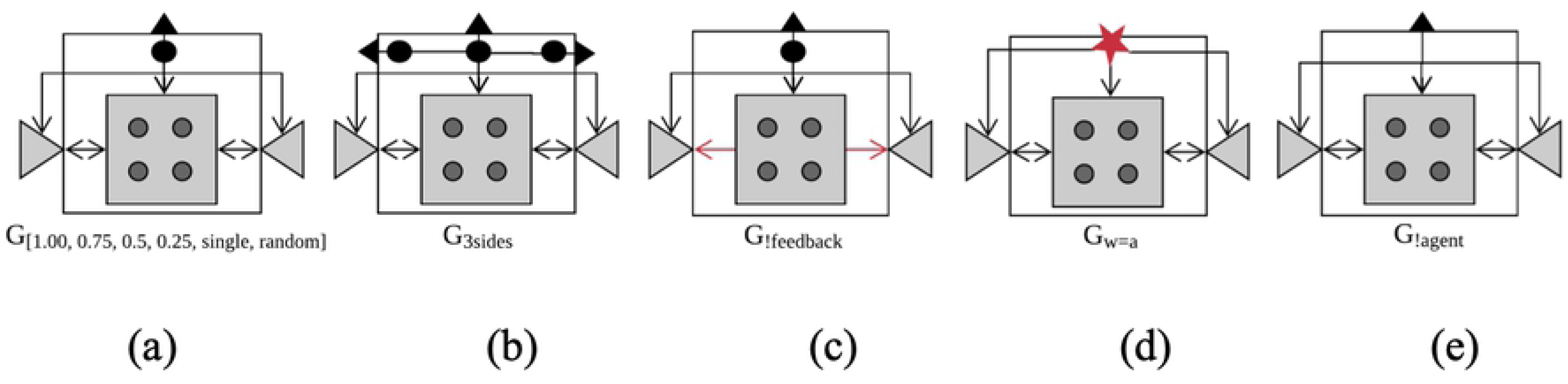
Schematic architecture of the five different animat designs. The animats have two motor units (grey triangles), four hidden units (dark grey circles) and one to six sensor units (black/red shapes). **(a)** Baseline design as in [17]. **(b)** Animat with sensors on three sides. There is an animat sensor and a wall sensor on each side. **(c)** Animat without feedback motors (motors cannot be part of the memory network). **(d)** Animat with a single sensor unit, measuring wall and animat simultaneously. **(e)** Animat without an animat sensor. Note that the architectures depict the maximal amount of units available. Whether any given unit is actually used depends on the evolved connectivity and logic function. Animats are initialized without connections between units.

### Design of the 2D Environment

All experiments simulated a two-dimensional environment. The world has ***32*32*** units (see Fig 16). All animats started on one of ***72*** predefined, uniformly distributed, starting positions. The selection for the starting position, as well as an animat’s initial orientation, was random. The original environment (see Fig 16(a)) had two rooms, which are connected by a gate. The animats’ goal was to travel between the two rooms in order to achieve a high fitness value. This design was adapted from the work of Koenig et al. [20]. As an additional dimension for evaluating reliability under environmental change, we tested all evolved MBs (the final generation) in additional environment designs (see Fig 16(b-e)).

**Fig 16.**
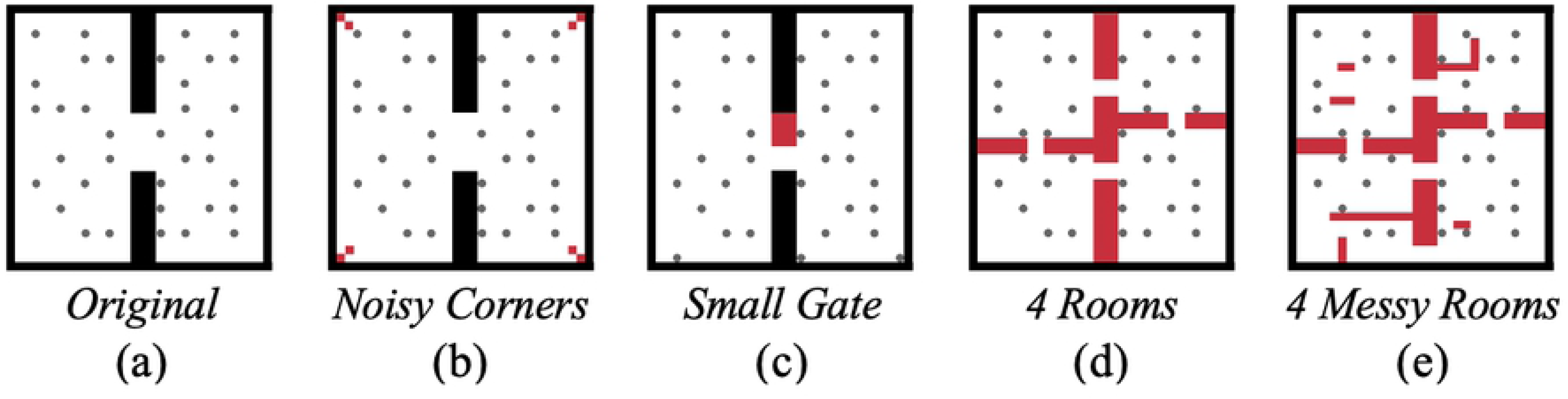
Environmental design. **(a)** The two-dimensional environment is based on a discrete grid architecture and contains two rooms. Animats draw a random starting position. Their orientation can be up, down, left, and right and is also randomly selected at initiation. **(b-e)** Four additional rooms were used to test the reliability of the animats. Red blocks mark the changes/additions in the room and represent walls. In (d), all four gates count as possible rewards. In (e), only gates on the vertical mid-line provide rewards.

We chose MBs as a simplified model of an artificial brain, since the basic idea of an MB is to emulate the recurrent connectivity structure found in real neural networks in a simple manner, while being complex enough to represent a cognitive system [15]. Furthermore, a recent study showed that MBs can be very compatible against variations of *artificial neural networks* and even showed higher performance in general [16]. Nevertheless, it would, in principle, also be possible to use a finite state machine [20], or artificial neural networks [30] to solve these kinds of tasks.

### Experiment Design

We selected ***G_0.50_*** to be the baseline setup for evolution, to which we compared all other evolutionary setups. This was because ***G_0.50_*** showed the highest reliability across group sizes. In sum, we came up with ***15*** different setups for the evolution of the animats. Using the MABE framework, we simulated each evolutionary setup ***30*** times. In each of these ***30*** evolutions, the EA had ***10,000*** generations to converge on the final solution. ***100*** genomes were mutated and evaluated in each generation. Each of these evaluations was repeated ***30*** times with different starting positions, orientation, and selection order (for the serial processing of the animats’ movement). After a genome was tested 30 times, it received a fitness score, which was computed based on the mean across the task performance of 30 single animats, with one being picked randomly from each of the ***30*** random test runs. In addition, in setup ***G_random_*** the group size varied for each of the ***30*** tests (drawn randomly from **72\***[***1, 0.95, 0.9, …, 0.1, 0.05, 0.01389***]).

### The Simulated Life

The fitness function that determines the probability of a genome being reproduced depends on two factors. First, animats have to travel as often as possible through the gate (change the room) (see Fig 16). Second, the animats need to avoid colliding with each other. Fischer et al. [17] showed the formal definitions of the fitness function as a weighted sum of the penalty for collision and the reward for crossing the gate. The weight of the reward (factor ***1.0***) is higher than the weight in the case of a penalty (factor ***0.075***). These weights need to be chosen carefully. If the penalty is too low or the reward is too high, animats will keep moving from one room to the other through the gate (herding effect) and ignore the penalty. On the other hand, given a high penalty and low reward, animats will evolve hardly any movement. To further reduce the herding effect around the gate, there is a refractory period of ***100*** timesteps after receiving a reward before an animat can receive another reward. Since each trial has a duration of ***500*** timesteps, any one animat can receive a total fitness score of at most ***4*** [17].

To further raise the task difficulty and to investigate the coordination and cooperation of animats in groups, we let animats co-exist in the same environment (in contrast to previous studies in this scope [15,18,21]). Currently, we have not implemented co-evolution and have only evaluated a genome by generating animats as identical clones (they have the same MB). There was no active knowledge exchange (“communication”) between animats in this study. Through the architecture of the animats, they have to develop the ability to distinguish which kind of sensory input to use for decision making. Sensors can only sense one position in front of (or on the side of) the animat and differentiate between static objects (walls) and dynamic objects (fellow animats), except for ***G_w=a_***.

Compared to the baseline setup, we included further control conditions in which animats did not receive the collision penalty and/or were not able to overlap. Those changes in the fitness function represented environmental rules which influenced the task difficulty. As a result, we were able to test dependencies between the evolution environment and the evolution of reliability.

### Post-Evolutional Evaluation

#### Reliability tests

The reliability tests were designed as follows: First, we selected the ***30*** genomes of generation ***10,000*** (***10k***) for each of the ***15*** conditions. Second, each genome was tested across ***21*** conditions varying in group size. To this end, we created groups of animat clones of the respective test group size for each of the ***30*15*** genomes. Test group sizes were uniformly distributed between ***1*** and ***72***. The interval of the relative distribution is [***0.0139, 0.05, 0.1, …, 0.9, 0.95, 1.0***]. A single animat is obviously not a group, but we treat it as one in order to simplify notation.

In addition to the reliability tests across varying group sizes in the baseline task design (*Original*), we created four modified test environments, as shown in Fig 16 (*Noisy Corners*, *Small Gate*, *4 Rooms*, *4 Messy Rooms*). Moreover, we included three additional test conditions in which we varied the interaction properties of the animats (*No Penalty*, *Blocked, Blocked and no penalty*). Finally, we tested each of the ***30*15*21*** different configurations in each of the eight test environments.

For the statistical analysis and the main reliability evaluations, we defined a reliability measure across group sizes in the *Original* environment design: R = ***〈TF〉_GS_***. The modified test environments represented four independent samples of possible environmental modifications and were only evaluated on their own for this reason. The results of the remaining three test conditions with varying interaction properties mainly served to highlight differences between the evolutionary setups, rather than testing reliability per se.

#### Brain complexity

To evaluate the complexity of the evolved MBs, we employed two complimentary measures provided by integrated information theory (IIT) [18, 43], ***Φ^Max^*** and the associated number concepts (#***Concepts***(***Φ^Max^***)). A major advantage of the measures developed within the IIT framework is that we can quantify the internal mechanisms (causal relations) of animats and their interactions (e.g., [44, 45]), which let us construct premises on how the cognitive processes work. The core of IIT’s measures is an information theoretic, and probabilistic graph analysis [18] based on the state-to-state transition probabilities of the units, i.e., their update functions. Please refer to [18, 19] for details on the evaluation. All calculations were conducted using the IIT Python package *pyphi* [23], which we used in our work to calculate ***Φ^Max^*** and the corresponding number of concepts. ***Φ^Max^*** represents the highest possible integrated information the system can achieve across all its subsets, which we used as an indicator for brain complexity. A concept is a set of physical mechanisms (e.g., neurons) that create integrated information [18]. Since the employed measures are state-dependent, we evaluated ***Φ^Max^*** and the number of concepts for every state a MB experienced during a lifetime (one trial) and selected the maximum value over all states as in [19]. Fig S1 (Supporting Information S2) shows by way of example that it is essential for high ***Φ^Max^*** in a system that many elements be integrated, meaning also maintaining feedback loops within the system. In this study, we only considered the brain complexity of the final generation (***10k***) due to the computational complexity of calculations using *pyphi*.

#### Statistics

The evolved fitness values, the reliability ***R***, and the IIT brain complexity measures were statistically evaluated across all evolutionary setups using a Kruskal-Wallis test, which showed a significant difference of the observed statistics between all groups taken together. Further, we used the Mann-Whitney-U test to evaluate the difference between pairs of evolutionary setups. Section S3 in the Supporting Information lists all statistical tests that are a subject of discussion in the results and discussion section.

---

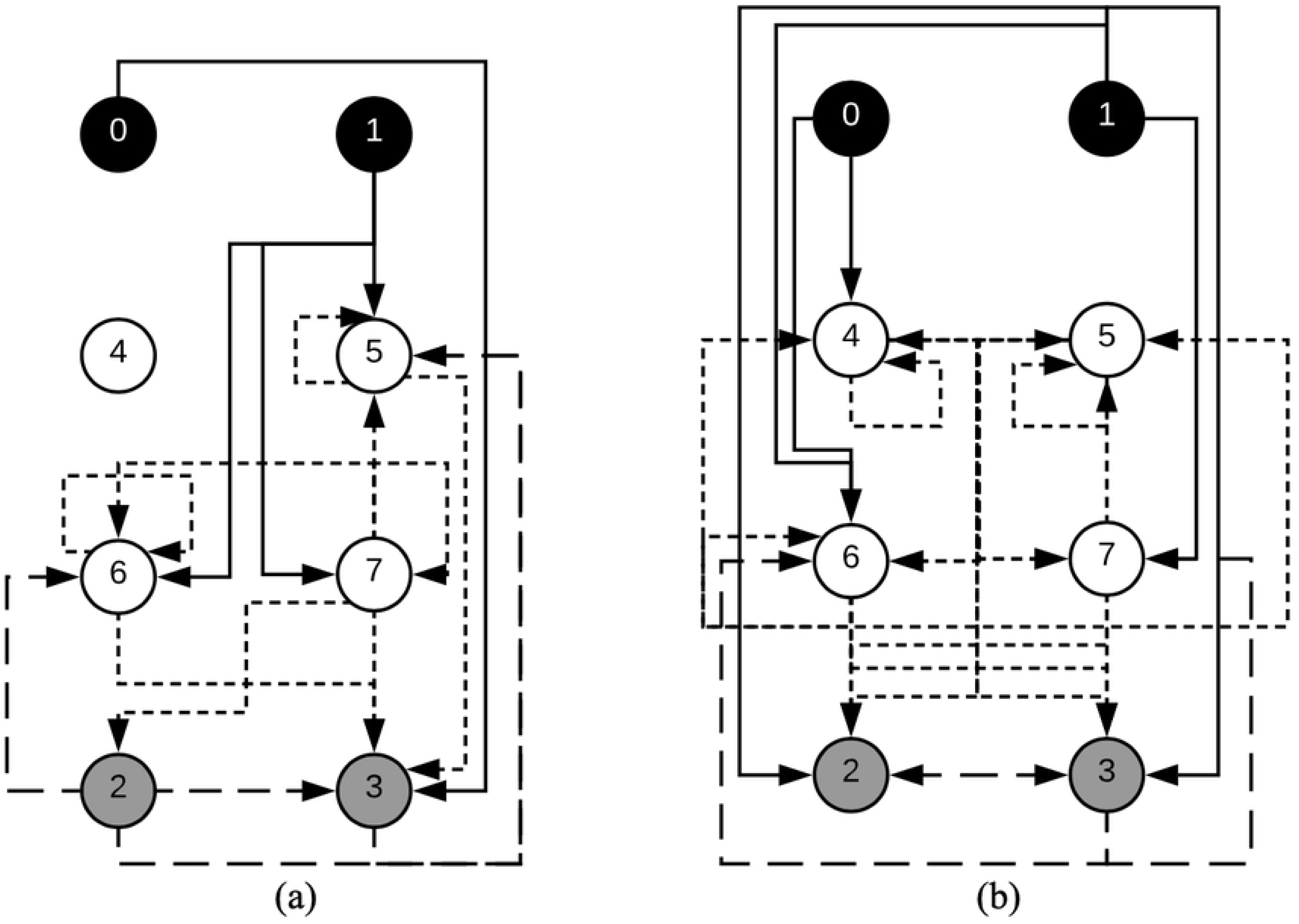

## References

1. Spearman C. “General Intelligence,” Objectively Determined and Measured. Am J Psychol. 1904;15: 201–292.

2. Gardner H. The theory of multiple intelligences. Ann Dyslexia. 1987;37: 19–35. doi:10.1007/BF02648057

3. Garnier S, Gautrais J, Theraulaz G. The biological principles of swarm intelligence. Swarm Intell. 2007;1: 3–31. doi:10.1007/s11721-007-0004-y

4. Pacala SW, Gordon DM, Godfray HCJ. Effects of social group size on information transfer and task allocation. Evol Ecol. 1996;10: 127–165. doi:10.1007/BF01241782

5. Dorigo M, Trianni V, Şahin E, Groß R, Labella TH, Baldassarre G, et al. Evolving Self-Organizing Behaviors for a Swarm-Bot. Auton Robots. 2004;17: 223–245. doi:10.1023/B:AURO.0000033973.24945.f3

6. Weick KE, Sutcliffe KM, Obstfeld D. Organizing for High Reliability: Process of Collective Mindfulness. Crisis Management. 2008. doi:10.1177/0020764009106599

7. Weick KE, Roberts KH. Collective Mind in Organizations: Heedful Interrelating on Flight Decks. Adm Sci Q. 1993;38: 357. doi:10.2307/2393372

8. Oliver N, Senturk M, Calvard TS, Potocnik K, Tomasella M. Collective Mindfulness, Resilience and Team Performance. Acad Manag Proc. 2017;2017: 12905. doi:10.5465/AMBPP.2017.12905abstract

9. Pinter-Wollman N, Penn A, Theraulaz G, Fiore SM. Interdisciplinary approaches for uncovering the impacts of architecture on collective behaviour. Philos Trans R Soc B Biol Sci. 2018;373. doi:10.1098/rstb.2017.0232

10. Engel D, Malone TW. Integrated information as a metric for group interaction. Dovrolis C, editor. PLoS One. 2018;13. doi:10.1371/journal.pone.0205335

11. List C, Philip Pettit. Group Agency and Supervenience. South J Philos. 2006;44: 1–22.

12. Walsh J, Ungson GR. Organizational Memory. Acad Manag Rev. 1991;16: 57–91. doi:10.2307/258607

13. Nonaka I. A firm as a knowledge-creating entity: a new perspective on the theory of the firm. Ind Corp Chang. 2000;9: 1–20. doi:10.1093/icc/9.1.1

14. Tsoukas H. The firm as a distributed knowledge system: A constructionist approach. Strateg Manag J. 1996;17: 11–25. doi:10.1002/smj.4250171104

15. Marstaller L, Hintze A, Adami C. The Evolution of Representation in Simple Cognitive Networks. Neural Comput. 2013;25: 2079–2107. doi:10.1162/NECO_a_00475

16. Hintze A, Kirkpatrick D, Adami C. The structure of evolved representations across different substrates for artificial intelligence. 2018; Available: http://arxiv.org/abs/1804.01660

17. Fischer D, Mostaghim S, Albantakis L. How swarm size during evolution impacts the behavior, generalizability, and brain complexity of animats performing a spatial navigation task. GECCO 2018. 2018; doi:10.1145/3205455.3205646

18. Oizumi M, Albantakis L, Tononi G. From the Phenomenology to the Mechanisms of Consciousness: Integrated Information Theory 3.0. Sporns O, editor. PLoS Comput Biol. 2014;10: 1–25. doi:10.1371/journal.pcbi.1003588

19. Albantakis L, Hintze A, Koch C, Adami C, Tononi G. Evolution of Integrated Causal Structures in Animats Exposed to Environments of Increasing Complexity. Polani D, editor. PLoS Comput Biol. 2014;10: e1003966. doi:10.1371/journal.pcbi.1003966

20. König L, Mostaghim S, Schmeck H. Decentralized evolution of robotic behavior using finite state machines. Hettiarachchi S, editor. Int J Intell Comput Cybern. 2009;2: 695–723. doi:10.1108/17563780911005845

21. Edlund JA, Chaumont N, Hintze A, Koch C, Tononi G, Adami C. Integrated Information Increases with Fitness in the Evolution of Animats. Graham LJ, editor. PLoS Comput Biol. 2011;7: e1002236. doi:10.1371/journal.pcbi.1002236

22. Marshall W, Gomez-Ramirez J, Tononi G. Integrated information and state differentiation. Front Psychol. 2016;7. doi:10.3389/fpsyg.2016.00926

23. Mayner WGP, Marshall W, Albantakis L, Findlay G, Marchman R, Tononi G. PyPhi: A toolbox for integrated information theory. Blackwell KT, editor. PLOS Comput Biol. 2018;14: e1006343. doi:10.1371/journal.pcbi.1006343

24. Miikkulainen R, Feasley E, Johnson L, Karpov I, Rajagopalan P, Rawal A, et al. Multiagent Learning through Neuroevolution. Lecture Notes in Computer Science (including subseries Lecture Notes in Artificial Intelligence and Lecture Notes in Bioinformatics). 2012. pp. 24–46. doi:10.1007/978-3-642-30687-7_2

25. Brown JL. Optimal group size in territorial animals. J Theor Biol. 1982;95: 793–810. doi:10.1016/0022-5193(82)90354-X

26. Olson RS. Elucidating the Evolutionary Origins of Collective Animal Behavior. PhD Proposal. 2015.

27. Olson RS, Hintze A, Dyer FC, Knoester DB, Adami C. Predator confusion is sufficient to evolve swarming behavior. J R Soc Interface. 2012;10: 20130305. doi:10.1098/rsif.2013.0305

28. Olson RS, Knoester DB, Adami C. Critical interplay between density-dependent predation and evolution of the selfish herd. Proceeding fifteenth Annu Conf Genet Evol Comput Conf - GECCO’13. 2013; 247. doi:10.1145/2463372.2463394

29. Karpov I V., Johnson LM, Miikkulainen R. Evaluating team behaviors constructed with human-guided machine learning. 2015 IEEE Conference on Computational Intelligence and Games (CIG). IEEE; 2015. pp. 292–298. doi:10.1109/CIG.2015.7317946

30. Stanley KO, Cornelius R, Miikkulainen R, Silva TD, Gold A. Real-time Learning in the NERO Video Game. Proc First Artif Intell Interact Digit Entertain Conf. 2005;2003: 2003–2004.

31. Stanley KO, Bryant BD, Miikkulainen R. Real-time neuroevolution in the NERO video game. IEEE Trans Evol Comput. 2005;9: 653–668. doi:10.1109/TEVC.2005.856210

32. Hamann H. Evolution of Collective Behaviors by Minimizing Surprise. 14th Int Conf Synth Simul Living Syst (ALIFE 2014). 2014; 344–351. doi:10.1145/2739482.2768497

33. Garnier S, Hamann H, Montes M, Christine DO, Eds TS, Hutchison D. Swarm Intelligence. In: Gerhard Goos, Hartmanis J, Leeuwen J van, editors. LNCS 8667. Brussels; 2014.

34. Ishiwata H, Noman N, Iba H. Emergence of Cooperation in a Bio-inspired Multi-agent System. Lecture Notes in Computer Science (including subseries Lecture Notes in Artificial Intelligence and Lecture Notes in Bioinformatics). 2010. pp. 364–374. doi:10.1007/978-3-642-17432-2_37

35. Reid CR, Lutz MJ, Powell S, Kao AB, Couzin ID, Garnier S. Army ants dynamically adjust living bridges in response to a cost-benefit trade-off. Proc Natl Acad Sci U S A. 2015;112: 15113–8. doi:10.1073/pnas.1512241112

36. Joshi NJ, Tononi G, Koch C. The Minimal Complexity of Adapting Agents Increases with Fitness. PLoS Comput Biol. 2013;9. doi:10.1371/journal.pcbi.1003111

37. Sheneman L, Hintze A. Evolving autonomous learning in cognitive networks. Sci Rep. Springer US; 2017; 1–11. doi:10.1038/s41598-017-16548-2

38. Beer RD, Williams PL. Information Processing and Dynamics in Minimally Cognitive Agents. Cogn Sci. 2015;39: 1–38. doi:10.1111/cogs.12142

39. Lizier JT, Prokopenko M, Zomaya AY. A Framework for the Local Information Dynamics of Distributed Computation in Complex Systems. 2014. pp. 115–158. doi:10.1007/978-3-642-53734-9_5

40. Zenil H. Compression-based investigation of the dynamical properties of cellular automata and other systems. Arxiv Prepr arXiv09104042. 2009; 1–25. Available: http://arxiv.org/abs/0910.4042

41. Clifford Bohm, Nitash C. G. AH. MABE (Modular Agent Based Evolver): A framework for digital evolution research. Proceedings of the European Conference on Artificial Life. MIT Press; 2017. pp. 76–83.

42. Hintze A, Edlund JA, Olson RS, Knoester DB, Schossau J, Albantakis L, et al. Markov Brains: A Technical Introduction. 2017; Available: http://arxiv.org/abs/1709.05601

43. Tononi G. Integrated information theory. Scholarpedia. 2015;10: 4164. doi:10.4249/scholarpedia.4164

44. Marshall W, Kim H, Walker SI, Tononi G, Albantakis L. How causal analysis can reveal autonomy in models of biological systems. Philos Trans R Soc A Math Phys Eng Sci. 2015; 1–22. doi:10.1098/not

45. Albantakis L. A Tale of Two Animats: What does it take to have goals? 2017;

